# Purkinje cell intrinsic activity shapes cerebellar development and function

**DOI:** 10.1101/2024.09.27.615345

**Authors:** Catarina Osório, Joshua J. White, Paula Torrents Solé, Nienke Mandemaker, Federico Olivero, Freya Kirwan, Fred de Winter, Eleonora Regolo, Francesca Romana Fiocchi, Inês Serra, Saffira Tjon, Zeliha Ozgur, Mirjam C.G.N. van den Hout, Wilfred F. J. van Ijcken, Guillermina López-Bendito, Aleksandra Badura, Lynette Lim, Geeske van Woerden, Martijn Schonewille

## Abstract

The emergence of functional cerebellar circuits is heavily influenced by activity-dependent processes. However, the role of intrinsic activity in Purkinje neurons, independent of external input, in driving cerebellar development remains less understood. Here, we demonstrate that before synaptic networks mature, Purkinje cell intrinsic activity is essential for regulating dendrite growth, establishing connections with cerebellar nuclei, and ensuring proper cerebellar function. Disrupting this activity during the postnatal period impairs motor function, with earlier disruptions causing more severe effects. Importantly, only disruptions during early development lead to pronounced defects in cellular morphology, highlighting key temporal windows for dendritic growth and maturation. Transcriptomic analysis revealed that early intrinsic activity drives the expression of activity-dependent genes, such as *Prkcg* and *Car8*, which are essential for dendritic growth. Our findings emphasize the importance of temporally-specific intrinsic activity in Purkinje cells for guiding cerebellar circuit development, providing a potential common mechanism underlying cerebellum-related disorders.

## Introduction

Neural development relies on a complex interplay between genetic and early activity-dependent processes that guide morphogenesis and establish neural circuits^1, 2^. Early electrical activity has been shown to regulate a wide range of developmental programs in various regions of the nervous system, including the retina^3, 4^, spinal cord^5, 6^, cochlea^7^, and neocortex^8, 9^. Genetic programs are especially relevant during cerebellar ontogenesis, where more than half of the brain’s neurons^10^ converge to form an intricate network of connections. The cerebellum undergoes a prolonged maturation period, making it particularly susceptible to developmental errors^11^. Indeed, evidence suggests that a common feature of several cerebellar-related brain disorders is the impairment of the Purkinje cell (PC) intrinsic activity^12^, which is the spontaneous activity generated by these principal neurons in the absence of synaptic inputs^13, 14^. However, the necessity of early PC activity for their maturation and its potential role in the development of cerebellar deficits remains largely unexplored. PCs are at the core of cerebellar circuitries, and their development is intertwined with the surrounding cerebellar elements^15^. The genetic and cellular programs that govern the genesis, migration, cell diversity, and differentiation of cerebellar cells have been explored and cataloged by recent single-cell transcriptomic analysis endeavors^16, 17^. Although less studied, activity-dependent mechanisms also play a crucial role in the early stages of cerebellar development. For example, the organization of PC parasagittal bands requires PC output signals^18^, while the competition and subsequent elimination of climbing fiber afferents depend on PC neuronal activity^19^. Additionally, electrical stimulation of cerebellar slices induces the expression of *Arc* in PCs, which contributes to the elimination of climbing fiber synapses^20^. Recent *in vitro* studies have shown that even when synaptic inputs are blocked, PCs exhibit intrinsic activity as early as postnatal day 3, with this activity gradually increasing until it stabilizes at mature levels by the end of the second postnatal week^21^. However, the extent to which early intrinsic activity influences the molecular programs underlying PC maturation, circuit formation, and overall cerebellar function remains unclear. To investigate the role of PC intrinsic activity in cerebellar development, we genetically reduced PC intrinsic activity at various postnatal stages in mice. Our findings reveal that early activity is essential for the development of PC dendritic arbors, axonal targeting of cerebellar nuclei cells, and the acquisition of cerebellar functions. Intrinsic electrical activity is most important during the first two postnatal weeks for PC maturation and for establishing balance and coordination functions. Molecularly, early intrinsic activity regulates the expression of several activity-dependent genes, such as *Prkcg* and *Car8*. Functional studies using shRNA knockdown of *Prkcg* and *Car8* demonstrate that they are key regulators of PC development. Overall, our results unveiled a selective requirement for early activity in shaping cerebellar development and function.

## Results

### The intrinsic activity of Purkinje cells controls morphological development

To reduce the intrinsic activity of PC during postnatal development, we crossed a tamoxifen-dependent *Pcp2^creERT2^* mice^22^ with a mouse line that conditionally expressed the inward rectifying potassium channel 2.1 (Kir2.1) fused to the mCherry reporter gene^23^ - *Pcp2^creERT2^;Kir2.1*. A control line *Pcp2^creERT2^;Ai14*^24^ was also generated, conditionally expressing only the tdTomato reporter gene. To examine the function of neuronal activity in PCs in a cell-autonomous manner, we administered a single low dose of tamoxifen (Tmx) at postnatal day 1(P1) to selectively express Kir2.1 in a subset of PCs in both control and mutant pups (Fig. 1a).

**Fig. 1:**
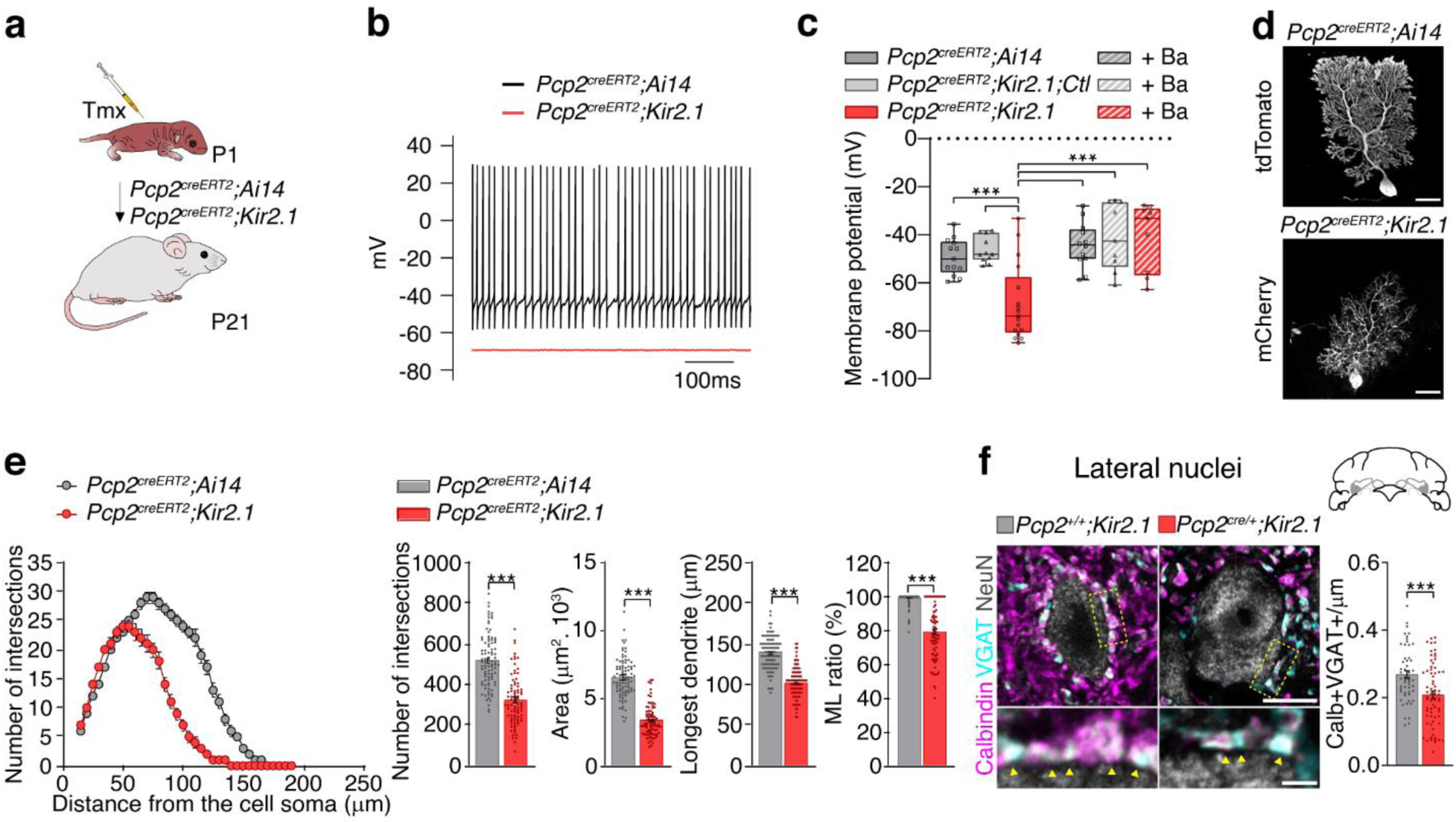
Decreased intrinsic activity impairs Purkinje cell development. **a**, Experimental design to reduce Purkinje cell activity by expressing Kir2.1. **b**, Example traces of spontaneous action potential firing recorded from *Pcp2^creERT2^;Ai14* (black) and *Pcp2^creERT2^;Kir2.1* (red) Purkinje cells at P21. **c**, Membrane potential in *Pcp2^creERT2^;Ai14* (n = 13 cells/3 mice), *Pcp2^creERT2^;Kir2.1,Ctl* (n = 11 cells/4 mice) and *Pcp2^creERT2^;Kir2.1* (n = 17 cells/4 mice) Purkinje cells, and the same groups in the presence of barium (300 µM Ba) (respectively n = 12 cells/5 mice, n = 9 cells/5 mice, n = 9 cells/5 mice). One-way ANOVA followed by Tukey’s multiple comparisons test \*\*\**P* < 0.001. **d**, Representative images of Purkinje cell morphology in *Pcp2^creERT2^;Ai14* and *Pcp2^creERT2^;Kir2.1* mice at P21. **e**, Sholl analysis, number of intersections, area, longest dendrite and ML ratio in *Pcp2^creERT2^;Ai14* (n = 92 cells/3 mice) and *Pcp2^creERT2^;Kir2.1* (n = 84 cells/3 mice) Purkinje cells. Unpaired *t*-test or Mann-Whitney *U* test \*\*\**P* < 0.001. **f**, Representative images (top) and insets (bottom) of Purkinje cell Calbindin-positive (+) axonal terminals (magenta) and presynaptic VGAT+ puncta (cyan) surrounding a NeuN+ neuron (grey) in the lateral nuclei of *Pcp2^+/+^;Kir2.1* and *Pcp2^cre/+^;Kir2.1* mice. Density of Calb+VGAT+ puncta contacting NeuN+ neurons in *Pcp2^+/+^;Kir2.1* (n = 56 cells/3 mice) and *Pcp2^cre/+^;Kir2.1* (n = 60 cells/3 mice) groups. Unpaired *t*-test with Welch’s correction \*\*\**P* < 0.001. Scale bar, 25 µm (**d**) Top = 10µm, Bottom = 2 µm (**f**). Bar data represent mean ± SEM. Tmx, tamoxifen

First, we confirmed the reduction of neuronal activity through *ex vivo* recordings from cerebellar slices. We compared PCs overexpressing Kir2.1 with two controls: PCs expressing only tdTomato (*Pcp2^creERT2^;Ai14*) and PCs that neither expressed Kir2.1 nor mCherry (*Pcp2^creERT2^;Kir2.1;Ctl*, where *Ctl* denotes control). Most control PCs fired spontaneously when no current was injected during current-clamp recordings. In contrast, Kir2.1 overexpression nearly abolished spontaneous action potentials in the majority of recorded PCs, significantly hyperpolarizing the resting membrane potential compared to control neurons. (Fig. 1b and 1c). As expected, applying 300µM barium, a potassium channel blocker, restored the membrane potential to control levels (Fig. 1c). Additionally, Kir2.1 overexpression caused a significant shift in the I/V curve of mutant PCs compared to controls, notably more hyperpolarized current at −140 mV. This I/V curve shift was also normalized in the presence of barium (Supplementary Fig. 1a, 1b, and 1c). Despite the loss of intrinsic activity, mutant PCs retained the ability to fire action potentials when positive current was injected (Supplementary Fig. 1d and 1e). These results demonstrate that PCs overexpressing Kir2.1 exhibit reduced intrinsic activity but maintain their capacity to generate action potentials when stimulated.

Next, we examined whether disrupting the intrinsic activity of PCs would affect their morphological development. We analyzed the dendritic arbors of PCs at P21 across different regions of the cerebellum. Sholl analyses revealed that PCs in *Pcp2^creERT2^;Kir2.1* mice had reduced dendritic complexity, smaller dendritic areas, shorter dendrite lengths, and lower occupancy in the molecular layer (ML) compared to control PCs. (Fig. 1e). These findings were corroborated by *in utero* electroporation (IUE) experiments (Supplementary Fig. 2).

To assess whether deficits in PC intrinsic activity affected presynaptic targeting as a population, we used two mouse lines, *Pcp2^+/+^;Kir2.1* (control) and *Pcp2^cre/+^;Kir2.1* (mutant). We evaluated the number of Calbindin-expressing (Calb+) and vesicular GABA transporter (VGAT+) boutons that contacted NeuN+ cerebellar nuclei cells. The reduction of PC intrinsic activity led to a significant decrease in the number of Calb+/VGAT+ contacts made with lateral NeuN+ neurons (Fig. 1f). Overall, these results indicate that reducing PC intrinsic activity during the first postnatal weeks disrupts dendritic growth and impairs presynaptic targeting.

### Disruption of the intrinsic activity of Purkinje cells impairs balance, coordination, and cerebellar-motor learning

Mutations that affect PC development are known to cause motor impairments^25–28^. Therefore, we investigated to what extent reducing the intrinsic activity of PCs from birth to adulthood could disrupt various aspects of motor behavior.

To test motor functions and cerebellar-dependent learning, we used adult (P60-P90) *Pcp2^cre/+^;Kir2.1* mice and control littermates, *Pcp2^+/+^;Kir2.1*. In the balance beam test, *Pcp2^cre/+^;Kir2.1* mice crossed the 12mm flat beam more slowly and made more missteps compared to controls (Fig. 2a, supplementary videos 1 and 2). These mutants were unable to cross a round 12mm beam (data not shown). Additionally, *Pcp2^cre/+^;Kir2.1* mice spent less time on the accelerating rotarod at both 40 and 80 rpm compared to controls (Fig. 2b and supplementary video 3). Using the LocoMouse test^29, 30^, *Pcp2^cre/+^;Kir2.1* mice exhibited abnormal body axis swings and a larger angle relative to the body axis (Fig. 2c). Although stride and stance distances were similar between groups, the mutants took significantly longer to complete these movements compared to controls (Supplementary Fig. 3b, supplementary video 4 and 5). Interlimb coordination was also disrupted in the mutants, with diagonal limb pairs not moving synchronously across different walking speeds (Supplementary Fig. 3c). However, in the open field test, while the distance traveled was similar between groups, the mutants moved faster than controls (Supplementary Fig. 3a). Taken together, our data revealed that overexpression of Kir2.1 in PCs from early development to adulthood results in an ataxic phenotype, characterized by impaired gait coordination and balance.

**Fig. 2:**
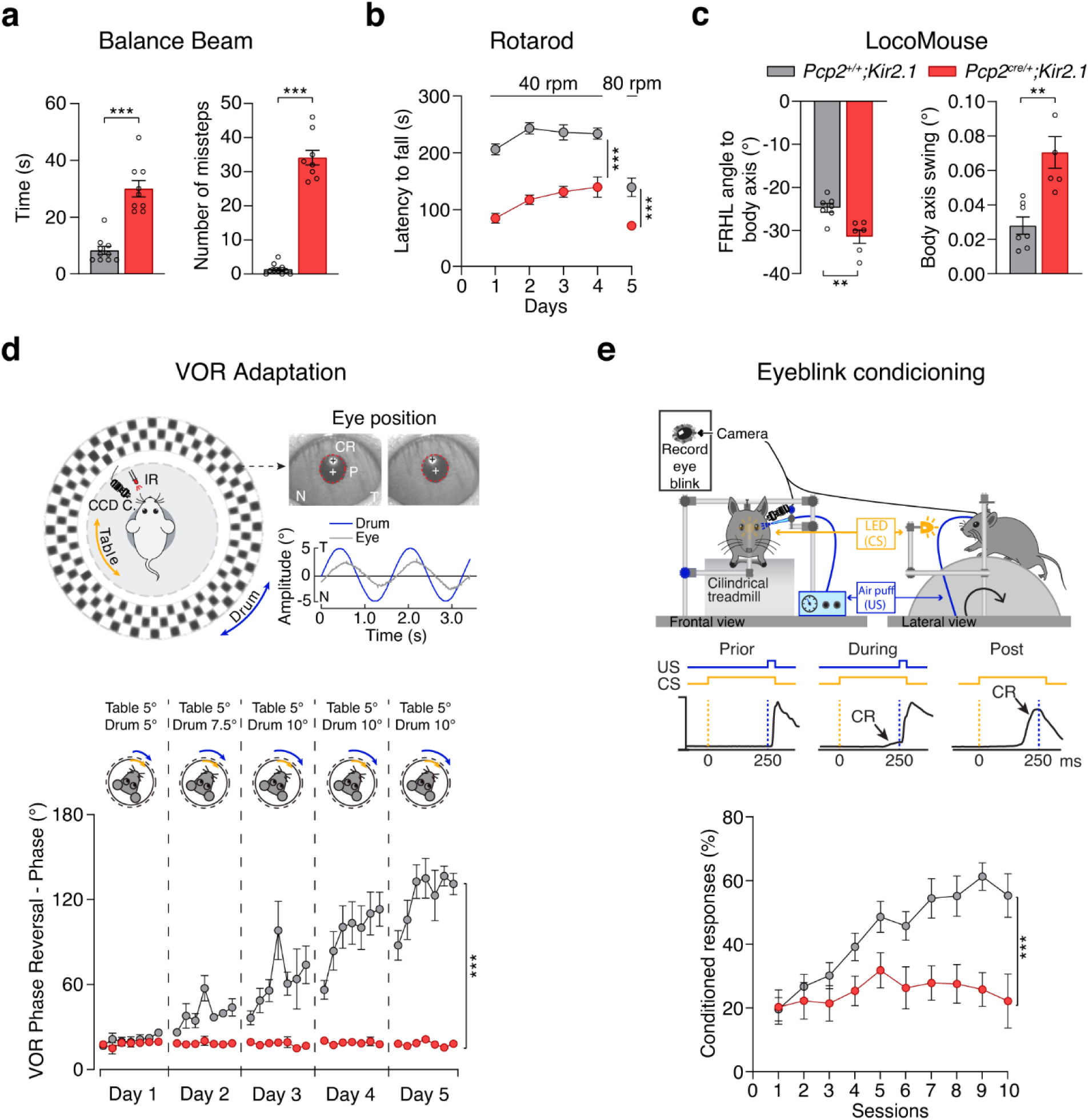
Reduced intrinsic activity of Purkinje cells impairs balance, coordination and the acquisition of cerebellar-motor learning. **a**, Balance beam test: Time to cross the beam and number of missteps made by *Pcp2^+/+^;Kir2.1* (n = 10 mice) and *Pcp2^cre/+^;Kir2.1* (n = 9 mice) groups. Mann-Whitney *U* test \*\*\**P* < 0.001. **b**, The latency to fall in the accelerated rotarod test in *Pcp2^+/+^;Kir2.1* (n = 11 mice) and *Pcp2^cre/+^;Kir2.1* (n = 10 mice) groups. Two-way ANOVA followed by Bonferroni’s multiple comparisons \*\*\**P* < 0.001. **c**, The angle of the front-right and hind-left (FRHL) paws diagonal to the central body axis and the angle of the body axis swing in *Pcp2^+/+^;Kir2.1* (n = 7 mice) and *Pcp2^cre/+^;Kir2.1* (n = 6 mice) groups. Unpaired *t*-test \*\**P* < 0.01. **d**, Illustration of compensatory eye movement recording setup. Turntable (yellow arrow) for vestibular stimulation and drum (blue arrow) for visual stimulation. CCD camera was used for infrared (IR) video-tracking of the left eye. N, nasal and T, temporal eye positions. Red circles = pupil fit; black cross = corneal reflection (CR); white cross = pupil (P) center. Example trace of eye position (gray) with drum position (blue). Quantification of vestibular-ocular reflex (VOR) phase reversal training in *Pcp2^+/+^;Kir2.1* (n = 7 mice) and *Pcp2^cre/+^;Kir2.1* (n = 7 mice) groups. Mixed-effect analysis \*\*\**P* < 0.001. **e**, Illustration of eyeblink conditioning recording setup. CS, conditioned stimulus (LED light - yellow); US, unconditioned stimulus (corneal air puff - blue). Percentage of conditioned responses in eyeblink conditioning training in *Pcp2^+/+^;Kir2.1* (n = 16 mice) and *Pcp2^cre/+^;Kir2.1* (n = 10 mice) groups. Two-way ANOVA \*\*\**P* < 0.001. Data represent mean ± SEM.

Given that PC activity is crucial for motor learning, we examined the impact of attenuated intrinsic activity on cerebellum-dependent learning behaviors, such as vestibular-ocular reflex (VOR) adaptation^31, 32^ and eyeblink conditioning (EBC)^33, 34^. We observed no differences in VOR and visual VOR gain (amplitude) and phase (timing) between the genotypes (Supplementary Fig. 3e and 3f). However, *Pcp2^cre/+^;Kir2.1* mutants exhibited lower gain and higher phase in the optokinetic reflex (OKR) compared to control (Supplementary Fig. 3d). In a phase-reversal VOR adaptation protocol, control animals successfully reversed their VOR direction by increasing phase over a 5-day experiment. In contrast, *Pcp2^cre/+^;Kir2.1* mutants failed to adapt their phase or gain, indicating impaired motor learning (Fig. 2d, Supplementary Fig. 3g). Similarly, in the EBC paradigm, where mice learn to associate a visual conditional stimulus (CS – LED light) with an eyeblink-inducing unconditional stimulus (US – air puff delivered to the mouse cornea), the mutant mice showed significantly reduced conditional response (CR – preventive eyelid closure) frequency and amplitude compared to controls (Fig. 2e, Supplementary Fig. 3h and 3i). Importantly, both genotypes had similar unconditional response peak times, confirming their ability to close the eyelid was intact (Supplementary Fig. 3j). In summary, these results demonstrated that reducing PC intrinsic activity impairs motor control and cerebellar-motor learning in this mouse model.

### The requirement of intrinsic activity for Purkinje cell morphological development decreases with age

After establishing the importance of PC intrinsic activity for dendritic growth and presynaptic targeting, we investigated whether this requirement persisted across different developmental stages. We administered low doses of Tmx at P1, P7, or P14 in *Pcp2^creERT2^;Ai14* and *Pcp2^creERT2^;Kir2.1* mice and analyzed PC dendritic arbor morphology at P7, P14, and P21, respectively (Fig. 3a, 3d and 3g). Following one week of Kir2.1 overexpression from P1 to P7, mutant PCs displayed more neurites emerging from the soma than controls (Fig. 3b and 3c). While control P7 PCs showed outward-oriented dendrites, those in PCs overexpressing Kir2.1 exhibited a more immature morphology, with neurites extending in all directions, resembling earlier developmental stages^35^. However, no significant differences were observed in the area or longest neurite length between mutant and control PCs (Fig. 3c). Overexpression of Kir2.1 during the second postnatal week led to a significant reduction in dendritic complexity, area, dendrite length, and occupancy in the ML in mutant PCs compared to controls at P14 (Fig. 3e and 3f). When intrinsic activity was reduced during the third postnatal week, mutant PCs at P21 still exhibited significant reductions in dendritic complexity, area, and dendrite length compared to controls (Fig. 3h, 3i). However, the extent of these reductions—10% in intersections, 21% in area, and 13% in longest dendrite length—was less pronounced compared to disruptions during the entire period of cerebellar development, P1 to P21 (38%, 49%, and 26%, respectively, Fig. 1d and 1e). Additionally, there was no significant difference in the percentage of dendritic occupancy in the ML between mutant and control PCs at this stage (Fig. 3i). Throughout these stages, we confirmed that Kir2.1 overexpression led to a reduced resting membrane potential and more hyperpolarized currents at −140 mV in mutant PCs at the various intermediate time points (Supplementary Fig. 4).

**Fig. 3:**
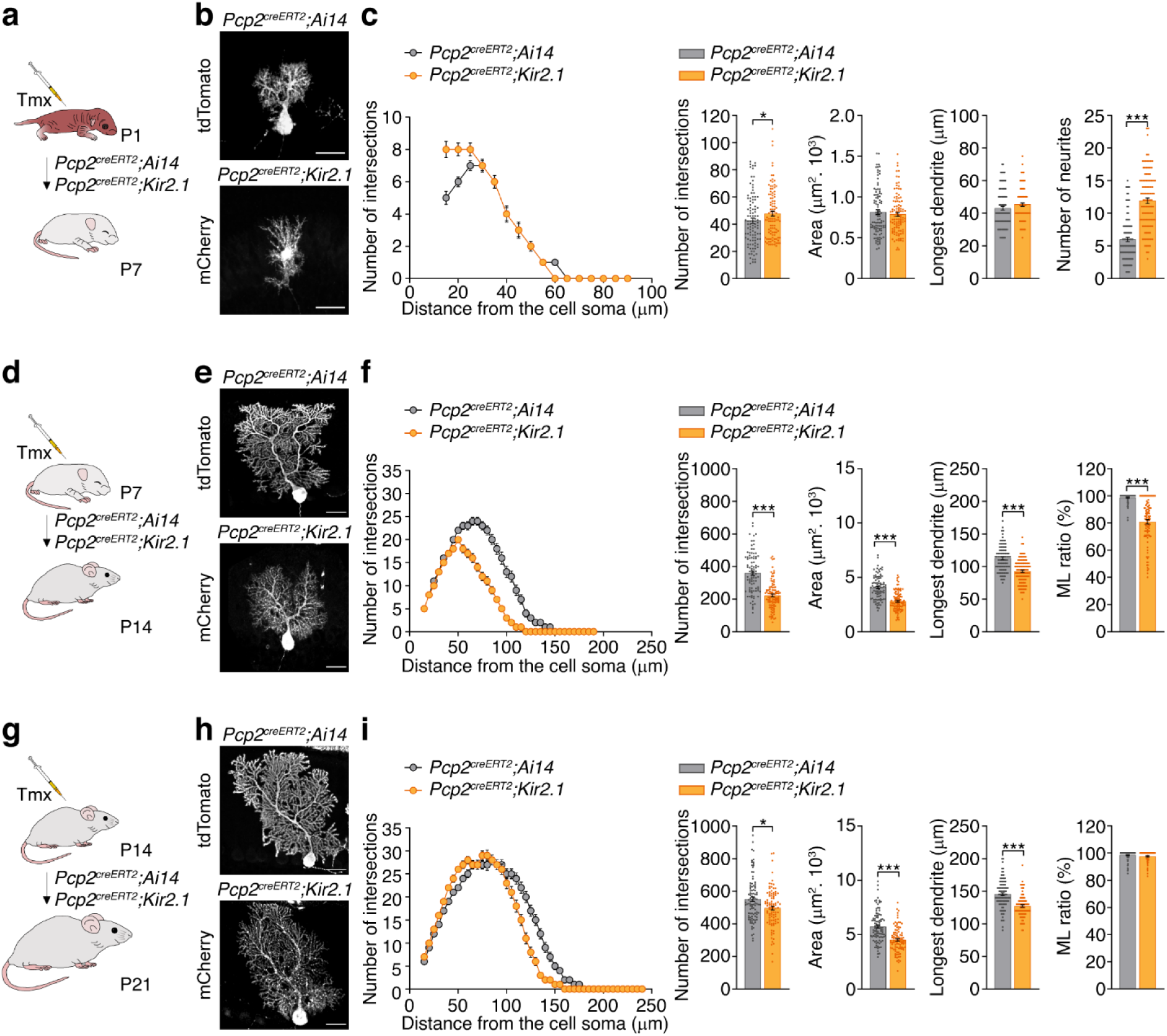
Decreased intrinsic activity delays Purkinje cell development in the first postnatal weeks. **a**, Experimental design to reduce Purkinje cell activity by expressing Kir2.1 from P1 to P7. **b**, Representative images of Purkinje cell morphology in *Pcp2^creERT2^;Ai14* and *Pcp2^creERT2^;Kir2.1* mice at P7. **c**, Sholl analysis, quantification of number of intersections, area, longest dendrite, and number of neurites in *Pcp2^creERT2^;Ai14* (n = 103 cells/3 mice) and *Pcp2^creERT2^;Kir2.1* (n = 104 cells/4 mice) Purkinje cells. Mann-Whitney *U* test \**P* < 0.05, \*\*\**P* < 0.001. **d**, Experimental design to reduce Purkinje cell activity by expressing Kir2.1 from P7 to P14. **e**, Representative images of Purkinje cell morphology in *Pcp2^creERT2^;Ai14* and *Pcp2^creERT2^;Kir2.1* mice at P14. **f**, Sholl analysis, quantification of number of intersections, area, longest dendrite and ML ratio in *Pcp2^creERT2^;Ai14* (n = 90 cells/3 mice) and *Pcp2^creERT2^;Kir2.1* (n = 101 cells/3 mice) Purkinje cells. Mann-Whitney *U* test and Unpaired *t*-test \*\*\**P* < 0.001. **g**, Experimental design to reduce Purkinje cell activity by expressing Kir2.1 from P14 to P21. **h**, Representative images of Purkinje cell morphology in *Pcp2^creERT2^;Ai14* and *Pcp2^creERT2^;Kir2.1* mice at P21. **i**, Sholl analysis, quantification of number of intersections, area, longest dendrite, and ML ratio in *Pcp2^creERT2^;Ai14* (n = 99 cells/3 mice) and *Pcp2^creERT2^;Kir2.1* (n = 95 cells/5 mice) Purkinje cells. Mann-Whitney *U* test and Unpaired *t*-test \**P* < 0.05, \*\*\**P* < 0.001. Scale bar, 25 µm (**b**, **e**, **h**) Data represent mean ± SEM. Tmx, tamoxifen

To assess how intrinsic activity affects PC connections with cerebellar nuclei cells at different developmental stages, we used *Pcp2^cre/+^;Kir2.1* mice from P1-P7 and *Pcp2^creERT2^;Kir2.1* mice from P7-P14 and P14-P21, administering high doses of Tmx to maximize Kir2.1 expression. Overexpression from P1-P7 significantly reduced the number of Calb+/VGAT+ contacts between PCs and NeuN+ neurons in the lateral cerebellar nuclei compared to controls (Supplementary Fig. 5a). Reducing activity from P7-P14 also significantly decreased these contacts (Supplementary Fig. 5b). However, no differences were observed in the number of inhibitory contacts when activity was reduced from P14-P21 (Supplementary Fig. 5c). These findings reveal that PC intrinsic activity is crucial for dendritic morphology and presynaptic targeting during development, with the most severe impairments occurring when activity is disrupted during the first two postnatal weeks.

### The onset of Purkinje cell intrinsic activity disruption determines the extent of motor impairment

We investigated whether the timing of intrinsic activity disruption in PCs during development influenced the severity of motor impairments. To test this, we administered high doses of Tmx at P7, P14 or P21 to maximize recombination of PCs expressing Kir2.1 in *Pcp2^creERT2^;Kir2.1* and their control littermates. We then assessed motor function in adulthood using the balance beam test in adulthood (Fig. 4a). Our findings showed that, regardless of when Tmx was administered, adult *Pcp2^creERT2^;Kir2.1* mice were significantly slower and made more missteps while crossing the flat beam compared to their respective controls, indicating impaired balance and coordination (Fig.4b and 4c). However, the severity of these motor impairments varied with the timing of activity reduction. Mice with disrupted PC activity starting from the second or third postnatal week performed better on the balance and coordination tests compared to those whose PC activity was reduced from birth (Supplementary Fig. 6). These results suggest that while intrinsic PC activity is crucial for optimizing motor movements, motor control is particularly sensitive to disruptions in PC activity during the first postnatal weeks.

**Fig. 4:**
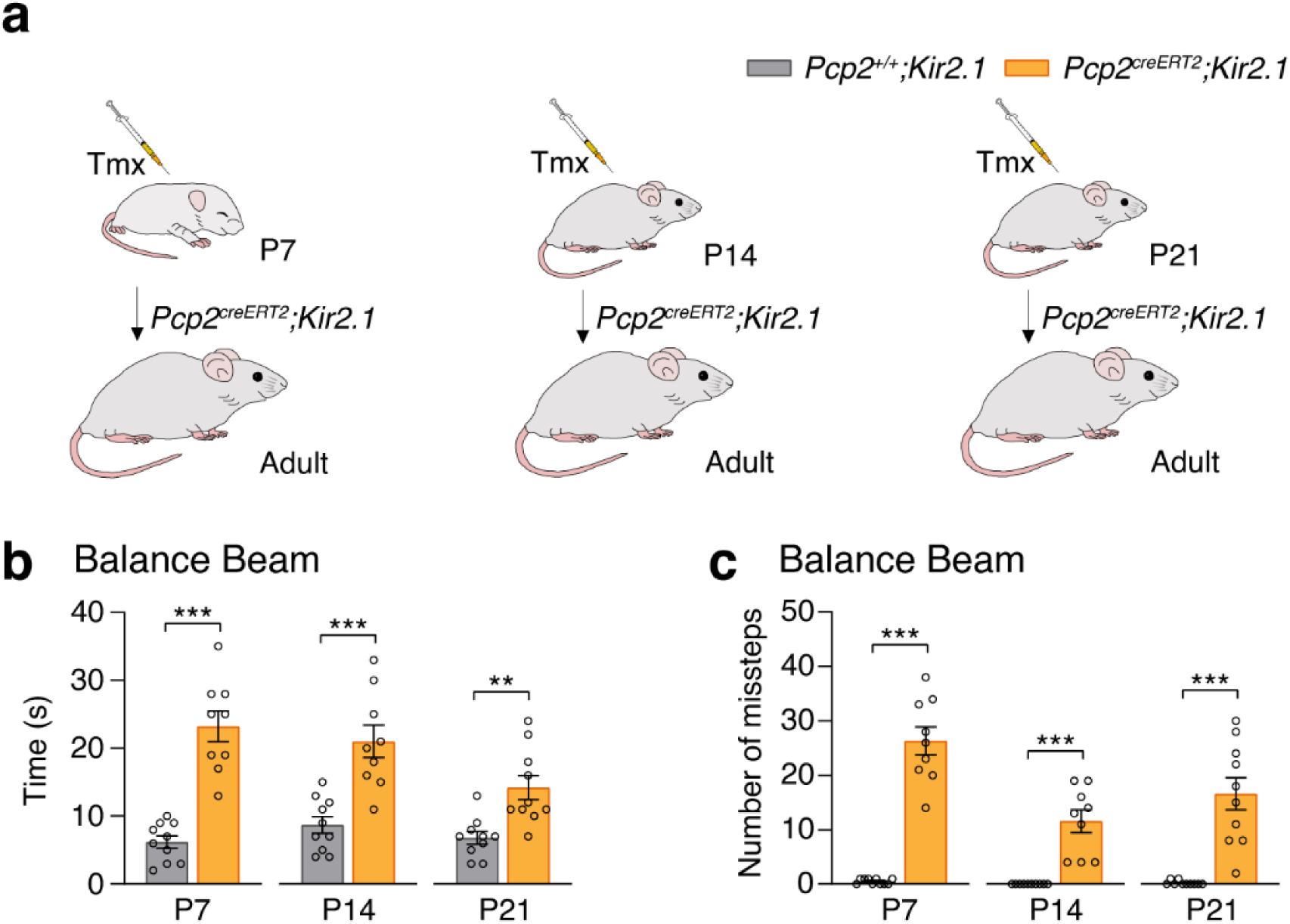
The onset of Purkinje cell activity reduction determines the severity of motor impairment. **a**, Experimental design to reduce Purkinje cell activity by expressing Kir2.1 from postnatal day 7 (P7), P14 or P21 to adulthood. **b**, Balance beam test: Time to cross the beam by *Pcp2^+/+^;Kir2.1* (n = 10 mice) and *Pcp2^creERT2^;Kir2.1* (n = 9 mice) groups. Mann-Whitney *U* test \*\*\**P* < 0.001. Unpaired *t*-test with Welch’s correction \*\**P* < 0.01, *** *P* < 0.001. **c**, Balance beam test: number of missteps made by *Pcp2^+/+^;Kir2.1* (n = 10 mice) and *Pcp2^creERT2^;Kir2.1* (n = 9 mice) groups. Mann-Whitney *U* test and Unpaired *t*-test with Welch’s correction *** *P* < 0.001. Data represent mean ± SEM. Tmx, tamoxifen

### Reduction of intrinsic activity in P7 Purkinje cells alters genes linked to synaptic development and movement disorders

To uncover the molecular mechanisms through which intrinsic activity influences PC development, we compared gene expression profiles between control PCs and *Pcp2^cre/+^;Kir2.1* PCs at P7. RNA sequencing (RNAseq) analysis identified 1,602 differentially expressed genes (DEGs) (Fig. 5a and Supplementary Fig. 7a, 7b and 7c). Gene ontology analysis revealed that many of these DEGs are involved in critical biological processes such as synaptic signaling and development (Fig. 7b). Enriched subgroups included “postsynaptic density membrane,” “presynaptic membrane,” “axon terminus,” and “transmembrane transport activity” (Supplementary Fig. 7d and 7e), which align with the significant morphological differences we observed. Given that disruptions in intrinsic PC activity lead to motor dysfunctions in our mouse model, we focused on DEGs associated with movement disorders. Using the KEGG Disease Database, we identified 216 genes linked to movement disorders with a cerebellar component, 25 of which were present in our DEG list (Fig. 5c and Supplementary Fig.7f and 7g). We selected two candidates for further investigation, *Prkcg* (protein kinase C gamma)^36^ and *Car8* (carbonic anhydrase VIII)^37^ (Fig. 5d), based on their implication in various forms of movement disorders^38, 39, 40^. Overall, our analysis indicates that reducing intrinsic activity in PCs disrupts the expression of genes crucial for synaptic signaling and development, as well as those associated with movement disorders.

**Fig. 5:**
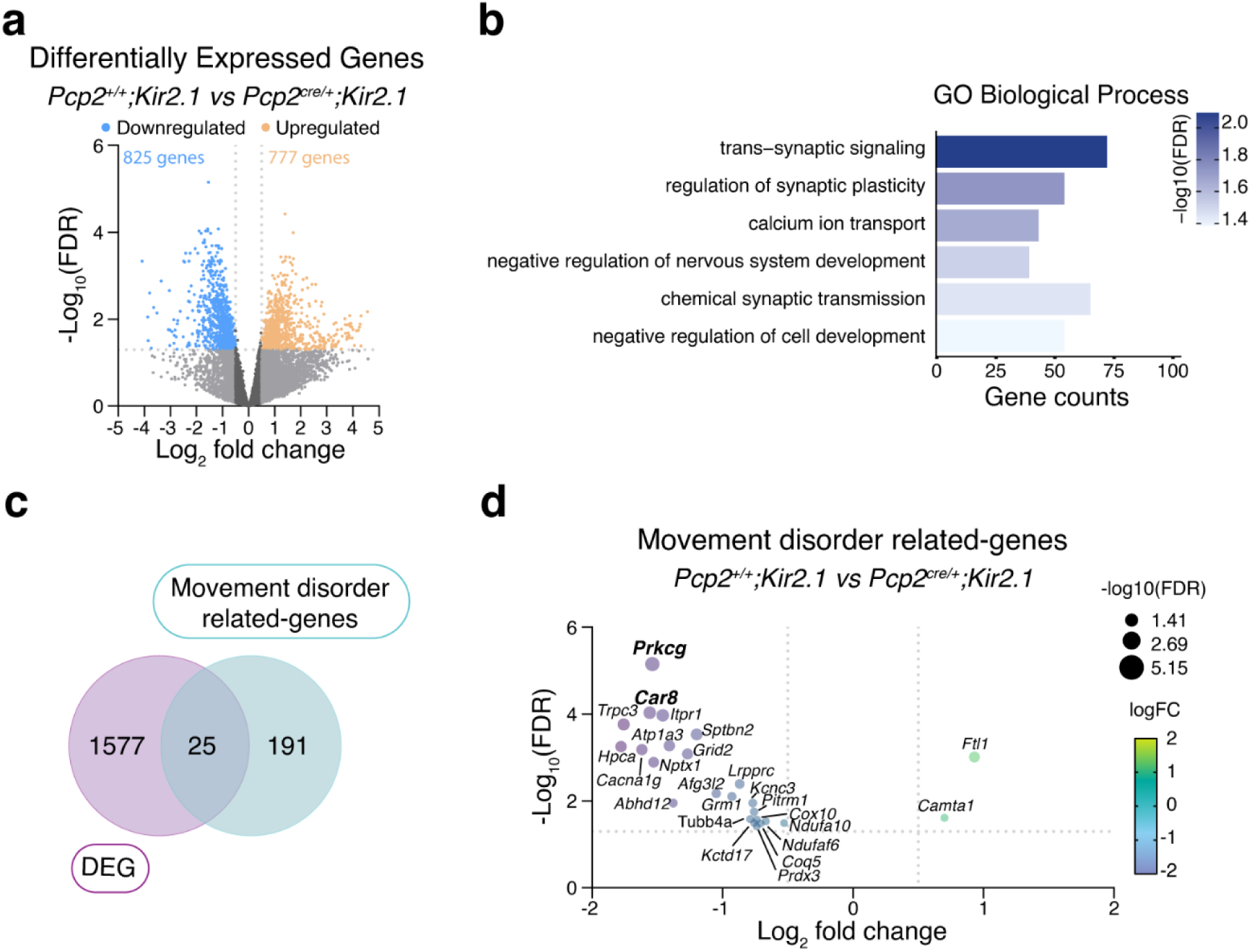
Decreased intrinsic activity of P7 Purkinje cells changes the expression of movement disorder-related genes. **a**, Vulcano plot representation of differentially expressed genes (DEG) between *Pcp2^+/+^;Kir2.1* and *Pcp2^cre/+^;Kir2.1* Purkinje cells at P7. Genes are significantly downregulated (blue) or upregulated (orange) within Log fold change > 0.5 and –Log_10_(FDR) > 1.3. **b**, Selected gene ontology (GO) enrichment analysis for the DEG biological process category plotted by gene count and –log_10_(FDR). **c**, Venn diagram showing the number of DEG at P7 in Purkinje cells when intrinsic activity is reduced, the number of genes associated with movement disorders with a cerebellar component from the KEGG Disease Database, and the 25 common genes between both lists. **d**, Vulcano plot illustrating the 25 common genes regarding their Log fold change and –Log_10_(FDR). FDR: false discovery rate.

### *Prkcg* and *Car8* regulate Purkinje cell dendritic development

To investigate whether the downregulation of *Prkcg* and *Car8* — genes associated with movement disorders — could be related to PC morphological development, we conducted *in vivo* gene knockdown experiments specific to PCs.^41^. Using effective short-hairpin RNAs (shRNAs) targeting *Prkcg* (*shPrkcg*) and *Car8* (*shCar8*), with *shLacZ* as a control (Supplementary Fig. 8a and 8c), we generated adeno-associated virus (AAVs) for each shRNA. These AAVs were injected into the lateral ventricles of P0 *Pcp2^cre/+^* mice, and tissue was collected at P7 for analysis (Fig. 6a). *In situ* hybridization confirmed the successful downregulation of *Prkcg* or *Car8* in mCherry-positive PCs (Supplementary Fig. 8b and 8d).

**Fig. 6:**
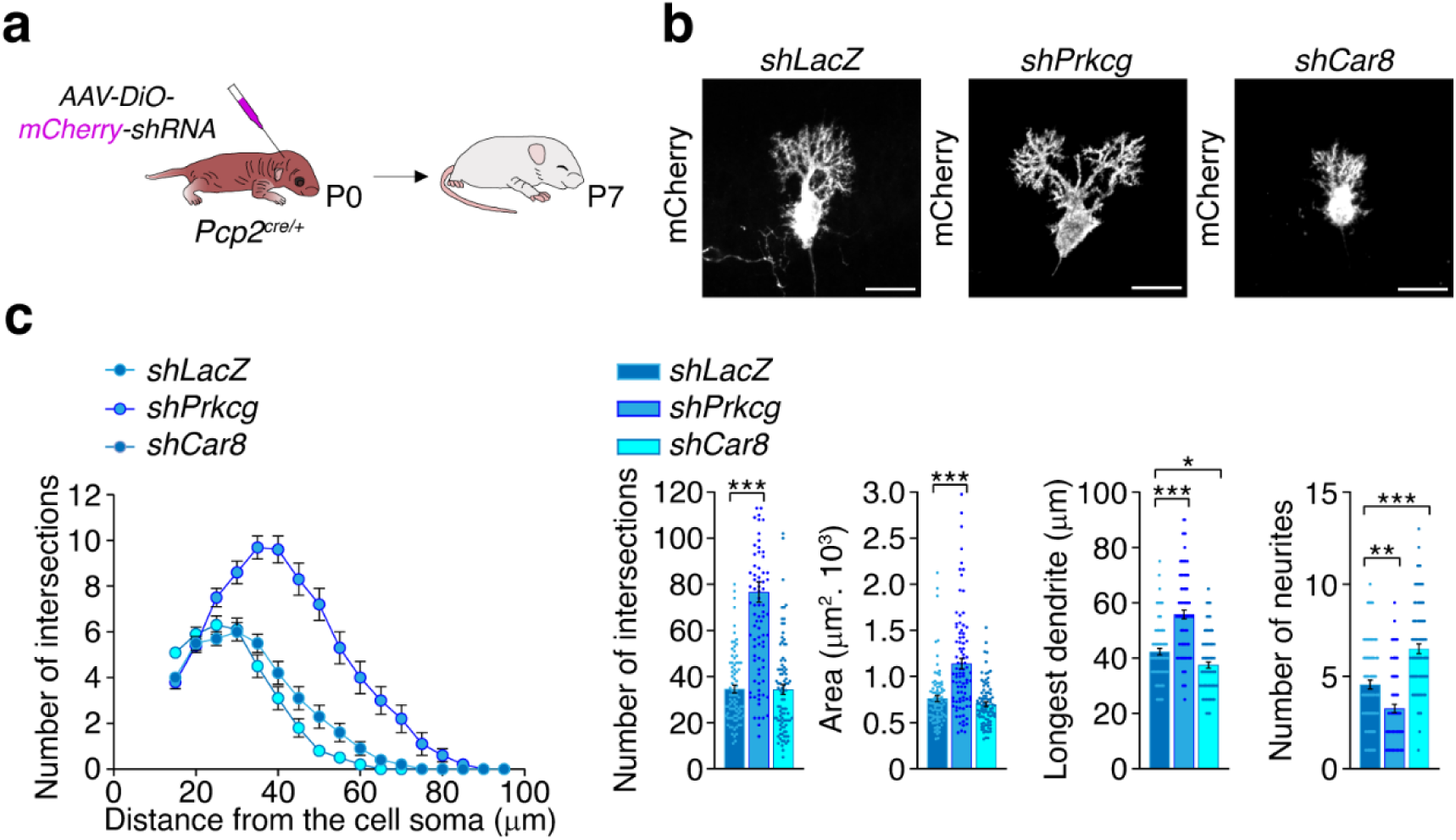
Activity-dependent *Prkcg* and *Car8* genes regulate Purkinje cell differentiation at P7. **a** Experimental design to knockdown gene expression in Purkinje cells using a shRNA viral vector. **b**, Representative images of Purkinje cell morphology in *shLacZ*, *shPrkcg,* and *shCar8* mice at P7. **c**, Sholl analysis, number of intersections, area, longest dendrite, and number of neurites in *shLacZ* (n = 84 cells/4 mice), *shPrkcg* (n = 90 cells/3 mice), and *shCar8* (n = 90 cells/3 mice) Purkinje cells. Kruskal-Wallis test/ANOVA followed by Dunn’s multiple comparisons \**P* < 0.05, \*\**P* < 0.01, \*\*\**P* < 0.001. Scale bar, 25 µm (**b**) Data represent mean ± SEM.

Morphological analyses revealed distinct effects on PC development when these genes were downregulated. Specifically, reducing *Prkcg* expression significantly increased dendritic complexity, the area of the dendritic arbor, and the length of the longest dendrite in P7 PCs, while also reducing the number of neurites emerging from the soma compared to controls (Fig. 6b and 6c). In contrast, downregulation of *Car8* did not affect dendritic complexity or overall arbor area. However, there was a reduction in the length of the longest dendrite and an unexpected increase in the number of neurites extending from the soma compared to control PCs, suggesting a delayed maturation in the knockdown cells (Fig. 6b and 6c). These findings indicate that at P7, intrinsic activity-dependent mechanisms regulate the expression of *Prkcg* and *Car8*, which play distinct roles in different aspects of PC dendritic maturation. Given that disruptions in these genes are associated with movement disorders, our results suggest that proper regulation of *Prkcg* and *Car8* is crucial for normal cerebellar function and motor control.

## Discussion

Early neuronal activity is known to regulate various aspects of development across multiple brain regions^2^, but the role of activity in the developing cerebellum remains poorly understood. Here, we report that postnatal growth, maturation, and function of PCs are heavily reliant on cell-intrinsic neuronal activity, also known as pacemaker activity. This activity is most crucial during the first two postnatal weeks, as disruptions during this period lead to profound effects on PC development. Genetic profiling revealed that early intrinsic PC activity drives the expression of numerous genes that regulate complex molecular pathways crucial for proper PC development. Disruptions in this activity resulted in significant alterations in these pathways, leading to impaired dendritic maturation.

Purkinje cells are featured by cell-intrinsic activity, which is commonly found to be affected in mouse models for various cerebellar disorders^12^. To explore the role early activity plays in neuronal development and neuronal circuit formation, we used a mouse model that allows for conditional overexpression of the Kir2.1 channel^23^, leading to hyperpolarization of neurons and, consequently, attenuation of neuronal excitability^8, 9, 42^. We found that PCs undergo activity-dependent differentiation and development, which is particularly relevant in the first two postnatal developmental weeks. Our results revealed that the reduction of neuronal activity results in an arrested development of the PC dendrite maturation. After one week of neuronal silencing, PC dendrite resembled what Ramón y Cajal has defined as “stellate cell with disoriented dendrites”^43^, characterized by the outgrowth of perisomatic protrusions from the cell body, unlike the control PCs in which it was possible to distinguish a small apical flattened dendrite coming from the soma. Moreover, reduction of activity in the second postnatal week of development led to defective morphology of the PC dendritic arbor compared with control. Similar effects on PC dendritic development have been observed in PCs of a mouse model deficient for Autism susceptibility candidate 2 (*Auts2)* gene^44^. *Auts2* regulates multiple developmental processes, and mutations in this gene are associated with autism spectrum disorders, intellectual disability, schizophrenia, and epilepsy^45^. Likewise, the number of PC VGAT terminals contacting the neurons in the cerebellar nuclei was reduced in conditions where PC activity was attenuated. The mechanisms underlying the reduction of PC contacts can be several and depend on the time during development the neuronal activity was initiated. For instance, failure of synapse formation, elimination of non-functional or less active synapses or/and lack of synapse maintenance have all been implicated in axonal targeting^46^. Additionally, it is possible that the development of the axons can be itself compromised. Our approach allowed for temporal control of the activity manipulation and consequently made it possible to determine the sensitive period. Interestingly, perturbation of activity only in the third postnatal week could not reproduce the severity of morphological defects observed after three weeks of neuronal silencing. Previous work has suggested that the first postnatal weeks in the mouse model are critical for most developmental processes that occur in the cerebellum^11^. For instance, delaying the expression of the mutant *Ataxin1* gene after cerebellar development is complete reduces disease severity, in particular, PC loss and impaired motor behavior^47, 48^. In a mouse model of autism, a PC-specific mutant for *Tsc1*, treatment with mechanistic target of rapamycin (mTOR) inhibitor, rapamycin, during cerebellar development, prevented PC death, the onset of impaired social behaviors in the adult mouse^49^ and restored the levels of intrinsic activity in the PC^50^. In a mouse model of spinocerebellar ataxia 1 (SCA1), the levels of intrinsic activity of PC are reduced, and restoring intrinsic activity using pharmacological intervention with 3,4-diaminopyridine reduces the locomotion phenotype^51^. Our data strengthen this concept by demonstrating that PC development during the first two postnatal weeks is particularly sensitive to changes in activity and that activity-dependent mechanisms are an integral part of PC maturation^52^.

Our results highlight the first two postnatal weeks as more sensitive to perturbations of PC activity. During this period, PCs are transiently connected with other PCs via axon collaterals, resulting in spontaneous network activation powered by PC intrinsic activity^52^, while the surrounding elements of the circuit are yet to become mature. Climbing fibers evolve from multi-innervating the PC soma to elimination, translation, and strengthening of the remaining synapses on the PC proximal dendrites^53^, the parallel fibers start innervating their distal dendrites^54^ while most of the molecular layer interneurons begin to integrate the nascent circuits^55^. The majority of PC-PC synapses have disappeared by the third postnatal week, while other synaptic inputs reach mature numbers and strength. Additionally, *in vitro* recordings of PC activity during development demonstrated that at the end of the second postanal week, the levels of intrinsic activity are close to those at mature ages and remain constant afterwards^21^. We posit that the immature cerebellum undergoes both morphological and physiological remodeling that relies heavily on early levels of PC intrinsic activity before the maturation of synaptic inputs. Disruption of early PC activity is sufficient to impair the emergence of a mature functional cerebellum.

Cook et al. have proposed that cerebellar disease is driven by PC intrinsic firing deficits preceding the onset of the first pathological phenotypes^12^. Indeed, a decrease in PC activity from birth led to motor dysfunction and abolishment of cerebellar-motor learning in our mouse model. Reduction of PC activity after the period of cerebellar development still resulted in motor impairments, but notably, the severity of motor deficiencies was reduced. Delaying the onset of PC dysfunction for one week after birth was sufficient to reduce the deficits observed in the balance beam test in adulthood, and this reduction was even more pronounced if PC were left unperturbed for two weeks after birth. Our findings align with deficits observed following repeated periods of hypoxia during the first two weeks, resulting in persistent cerebellar deficits^56, 57^). However, it remains to be determined whether early perturbation of PC intrinsic activity is sufficient to cause specific cognitive and emotional deficits, as previously demonstrated for later developmental periods^58^.

Early non-evoked spontaneous activity has been shown to be crucial to the development of many neuronal circuitries, including the retina, the spinal cord, the cochlea, and different regions in the neocortex^2, 59^. In particular, this early form of activity regulates several aspects of circuitry formation, such as neurogenesis, migration, apoptosis, and differentiation^60^. Perturbations of this early activity, whether due to prenatal drug exposure, viral infection, or perinatal hypoxia, have been linked to neurological disorders like epilepsy and autism spectrum disorders^60^. In the cerebellum, spontaneous, synchronized network activity, in the form of traveling waves, is known to be present in PCs in the first postnatal week of development, but this is transient and, therefore, not present in the adult cerebellum^52^. Here, we found that motor skills such as balance and coordination rely on the establishment of precise nascent circuits during cerebellum development.

Over the last few years, single-cell RNA sequencing and spatial mapping have unraveled genetic and cellular developmental programs in the mammalian cerebellum^16, 17^. Here, we explored the transcriptional programs underlying the activity-dependent maturation of PCs at the end of the first postnatal week. Our RNA sequencing data suggests that PC intrinsic activity induces a molecular program that regulates trans-synaptic signaling, cell development, and calcium transport. Indeed, silencing of activity at P7 reduced the number of contacts PC axons make with cerebellar nuclei cells and delayed PC development. Moreover, genes involved in calcium signaling (*Cacna1g*, *Grm1*, *Hpca*, *Trpc3*, *Camta1*, *Car8*, *Itpr1*, *Prkcg*) and mitochondrial function (*Ndufa10*, *Prdx3*, *Lrpprc*, *Afg3l2*, *Coq5*, *Pitrm1*, *Cox10*, *Nduf6*) were altered in their expression. Prior to the formation of synaptic networks, calcium signaling appears to be the primary source of activity regulating different aspects of neuronal development, including dendritic growth and patterning, axonal growth and pathfinding, and neurotransmitter specification^61^. Importantly, disruption of calcium homeostasis is a hallmark of many movement disorders related to cerebellum^12, 62, 63^, and such disruptions are known to impair PC intrinsic firing^64^. In accordance with this and based on our mouse model motor abnormalities, we sort our DEG based on movement disorders. We focused on *Prkcg* and *Car8*, two genes associated with movement disorders that also showed altered expression in our model. Our work extends these findings by directly linking their expression to neuronal activity, particularly during the critical period of PC maturation. *Prkcg* encodes for the protein kinase C gamma (PKCγ), serine/threonine kinase highly expressed in PCs^65^ and responsible for many neuronal functions such as synaptic maturation^66^, and regulation of long-term depression^67^. Mutations in the *Prkcg* gene are the cause of Spinocerebellar ataxia 14 (SCA14), characterized by gait imbalance, dysarthria, and abnormal eye movements^38, 68^. The pathology in SCA14 can have different underlying mechanisms depending on the mutation and type of PKCγ dysregulation. Both gain-of-function^69, 70^ and loss-of-function^71^ kinase activity have been reported in SCA14^72^. Interestingly, *in vitro* studies have shown that activation of PKCγ causes an inhibition of PC dendritic growth while inhibition of PKCγ promotes dendritic growth^73^. These observations are in accordance with our functional analysis *in vivo*, where the downregulation of *Prkcg* led to increased dendritic complexity, which contrasts with the stunted dendritic growth seen in PC models overexpressing the Kir2.1 channel. These data suggest that *Prkcg* may function as a suppressor of excessive dendritic growth branching during normal development, a role that becomes dysregulated in SCA14. The carbonic anhydrase-related protein 8 (CAR8) is an acatalytic member of the carbonate dehydratase family, and it is highly expressed in PCs^37^. *Car8* mutations have been described as the cause of cerebellar ataxia with intellectual disability and disequilibrium syndrome^39, 74^. Loss-of-function mouse models for CAR8 have ataxia and dystonia, and while they do not display gross morphological defects in adulthood, transient anatomical alterations are known to precede motor dysfunction^75, 76^. In our study, Car8 downregulation mirrored the abnormal dendritic development observed in the Kir2.1 overexpression model, highlighting the role of CAR8 in early Purkinje cell maturation and suggesting its involvement in reducing lateral processes and defining major dendritic branches. Many other DEGs identified in this work are likely involved in different aspects of PC maturation, but further research is needed.

In summary, our study provides compelling evidence that PC intrinsic activity plays a pivotal role in the early development of the cerebellum. By activating molecular programs that govern PC maturation, this activity ensures the proper formation of cerebellar circuits. Disrupting this activity early in development impairs the emergence of a mature, functional cerebellum and results in motor deficits later in life.

## Data availability

Sequencing data will be deposited at the National Center for Biotechnology Information BioProjects Gene Expression Omnibus (GEO) and will be available upon publication. All constructs generated in this study will be available upon publication.

## Acknowledgements

The authors kindly thank all personnel of the animal facility of the Erasmus Medical Center Rotterdam, Elize Haasdijk, Erika Sabel-Goedknegt, Laura Post and Doriane Gisquet for excellent technical assistance and Guillermina López-Bendito for providing the Kir2.1 (Gt(ROSA)26Sor^tm2(CAG-KCNJ2/mCherry)Fmr^) mouse model. This work was supported by H2020 European Research Council (ERC-Stg #680235) (M.S.), Dutch Organization for Life Sciences (ZonMw Off-road (ZonMW-451001027), OCENW.XS5.121 and Incentive Grant for Women in STEM (19498)) (C.O.), Dutch Organization for Life Sciences (VENI fellowship (NWO-ENW 016.Veni.192.270) and OCENW.XS21.1.087) (J.J.W.) and AAV vector production was supported by grants from KNAW Research fund and Leiden Regenerative Medicine Platform (FdW).

## Author contributions

Conceptualization – C.O., L.L., G.vW. and M.S.

Methodology – C.O. and J.J.W.

Investigation – C.O., J.J.W., P.T.S., N.M., F.O., F.K. and E.R.

Data curation – C.O., J.J.W., P.T.S., N.M., F.O., F.K., I.S. and S.T.

Formal analysis – C.O., J.J.W., P.T.S., F.R.F. and I.S.

Resources – C.O., J.J.W., F.dW., F.R.F., S.T., Z.O., M.C.G.N.H., W.F.J., G.L.B. and A.B.

Software – C.O., J.J.W., F.R.F. and S.T.

Visualization – C.O.

Writing – original draft – C.O.

Writing – review & editing – all authors

Funding acquisition – C.O., J.J.W., F.dW. and M.S.

Supervision – C.O., J.J.W., and M.S.

---

**Supplementary Fig. 1:**
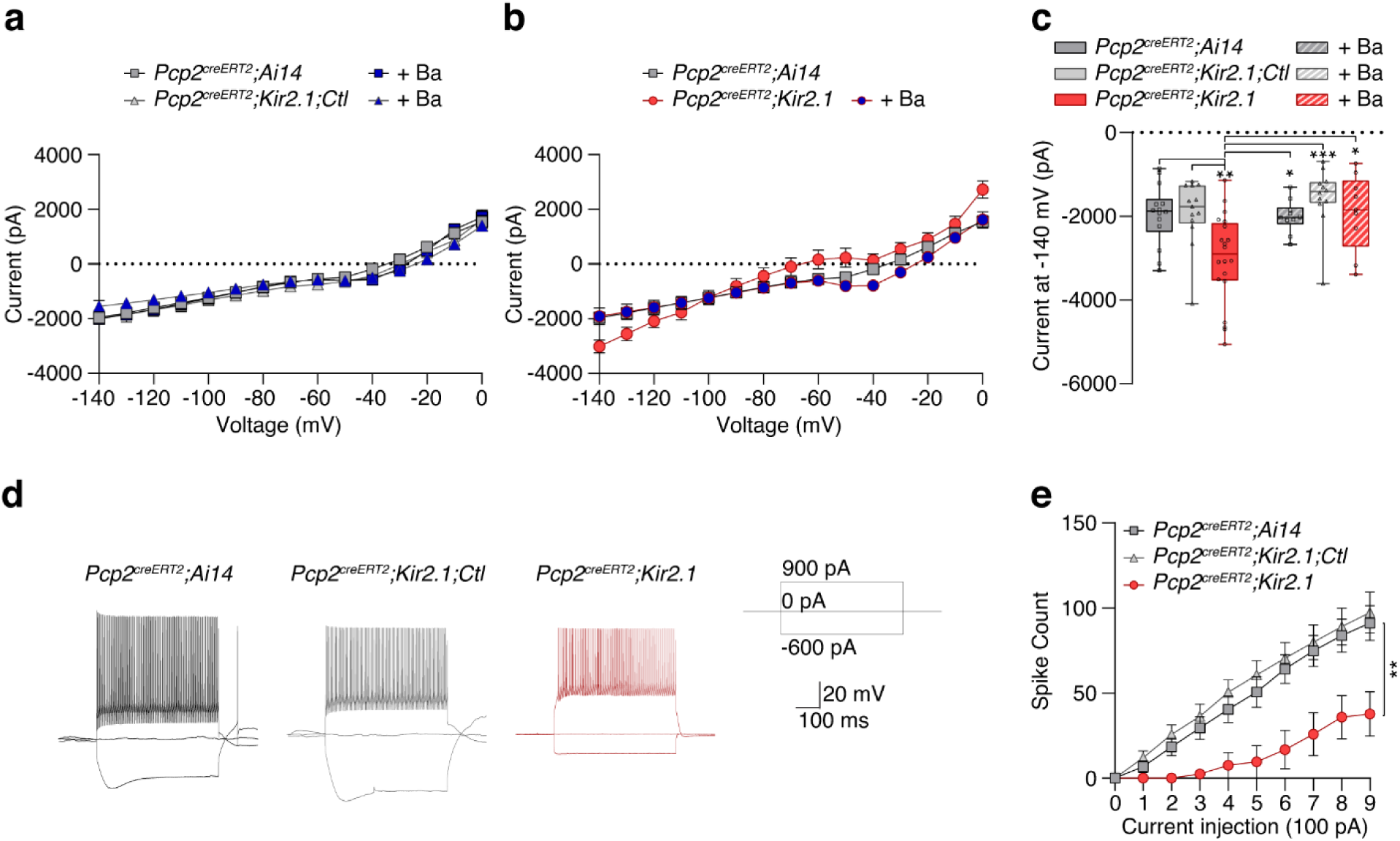
Overexpression of Kir2.1 decreases the excitability of Purkinje cells. **a**, Current-voltage relationship (I-V curve) in *Pcp2^creERT2^;Ai14* (n = 14 cells/4 mice) and *Pcp2^creERT2^;Kir2.1,Ctl* (n = 13 cells/5 mice) Purkinje cells under normal conditions and in the presence of barium at P21, respectively (n = 10 cells/5 mice; n = 12 cells/4 mice). **b**, Current-voltage relationship (I-V curve) in *Pcp2^creERT2^;Ai14* (n = 14 cells/4 mice) and *Pcp2^creERT2^;Kir2.1* (n = 20 cells/6 mice) Purkinje cells under normal conditions and in the presence of barium (n = 9 cells/3 mice). **c**, Quantification of the current amplitude in the absence or presence of barium for different groups hyperpolarized at −;140mV. One-way ANOVA followed by Tukey’s multiple comparisons test \**P* < 0.05, \*\**P* < 0.01, \*\*\**P* < 0.001. **d**, Representative whole-cell recordings obtained from *Pcp2^creERT2^;Ai14* (n = 13 cells/5 mice), *Pcp2^creERT2^;Kir2.1,Ctl* (n = 10 cells/4 mice) and *Pcp2^creERT2^;Kir2.1* (n = 9 cells/5 mice) Purkinje cells at P21 in response to current injections. **e**, Spike count of Purkinje cells from different groups in response to a series of current injections with 100pA increments. Two-way ANOVA followed by Tukey’s multiple comparisons test \*\**P* < 0.01.

**Supplementary Fig. 2:**
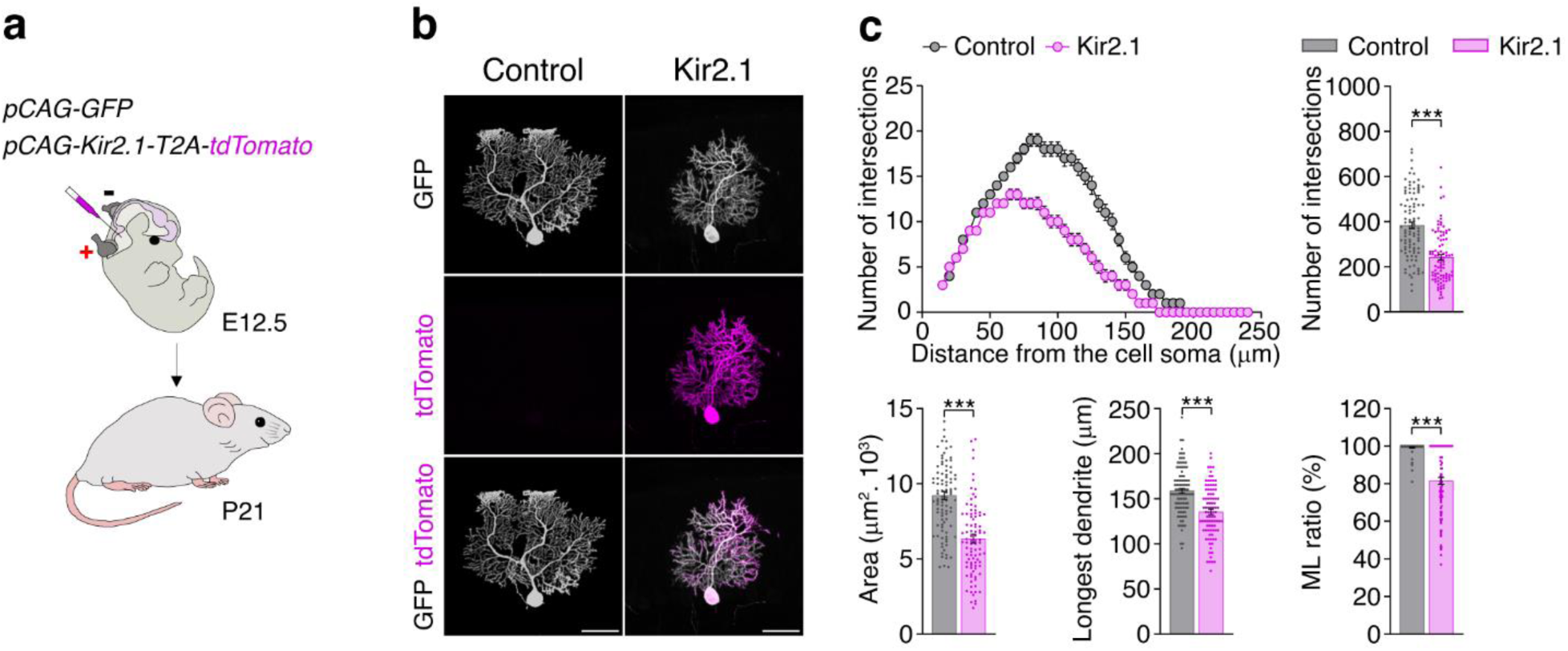
Overexpression of Kir2.1 impairs the development of Purkinje cell dendritic arbor. **a**, Experimental design to reduce Purkinje cell activity by expressing Kir2.1 in Purkinje cell progenitors using *in utero* electroporation on embryonic day 12.5 (E12.5). **b**, Representative images of Purkinje cell morphology in control (GFP) and Kir2.1 (tdTomato) at P21. **c**, Sholl analysis, number of intersections, area, longest dendrite, and ML ratio in control (n= 98 cells/4 mice) and Kir2.1 (n = 92 cells/4 mice) Purkinje cells. Mann-Whitney *U* test and Unpaired *t*-test \*\*\**P* < 0.001. Scale bar, 50 µm (**b**) Data represent mean ± SEM.

**Supplementary Fig. 3:**
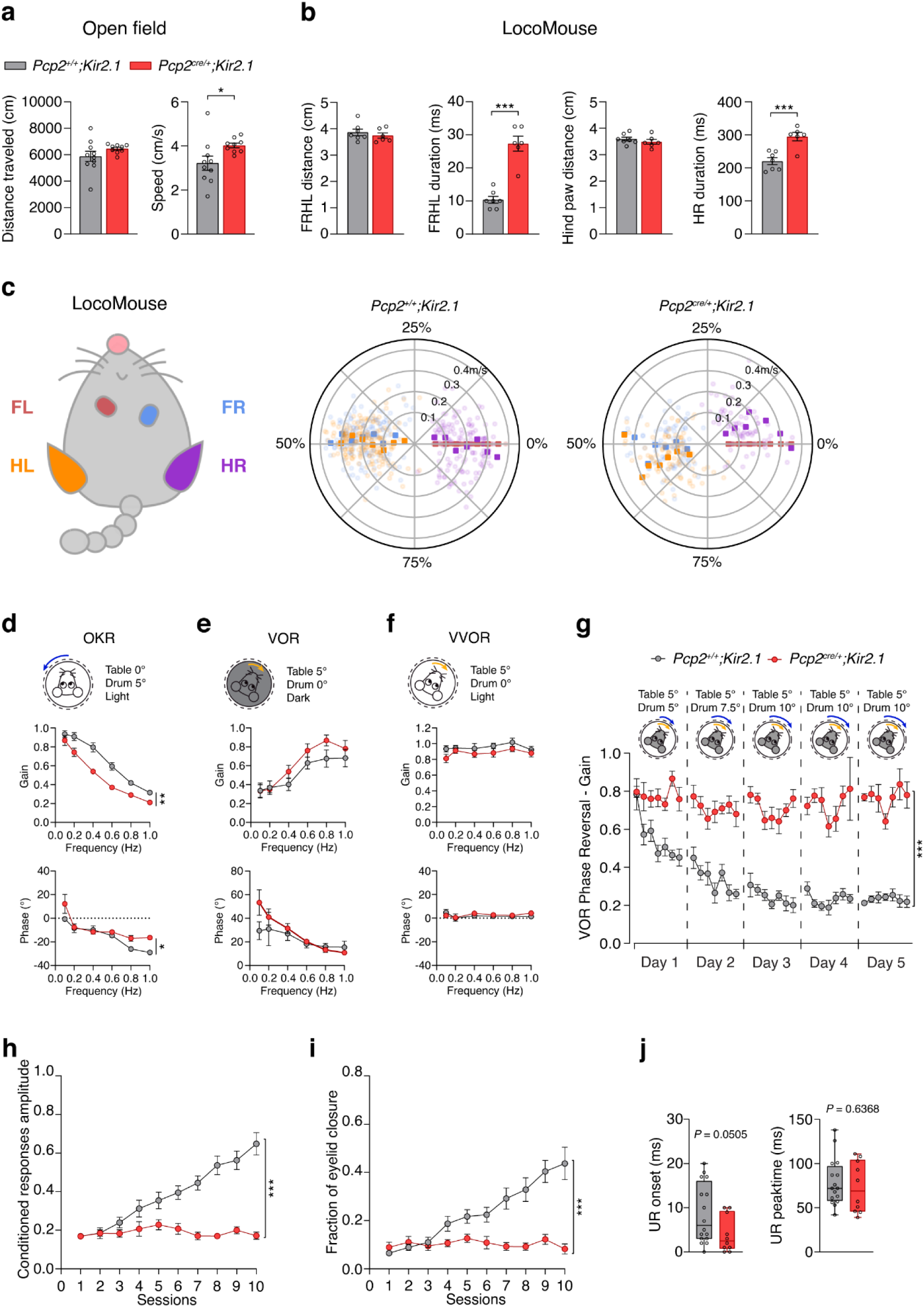
Reduction of Purkinje cell intrinsic activity impairs motor performance and motor learning. **a**, Open field test: Distance traveled and speed made by *Pcp2^+/+^;Kir2.1* (n = 10 mice) and *Pcp2^cre/+^;Kir2.1* (n = 9 mice) groups. Unpaired *t*-test with Welch’s correction \**P* < 0.05. **b**, LocoMouse test: FRHL distance, FRHL duration, hind paw distance, and HR duration **in** *Pcp2^+/+^;Kir2.1* (n = 7 mice) and *Pcp2^cre/+^;Kir2.1* (n = 6 mice) groups. Unpaired *t*-test \*\*\**P* < 0.001. **c**, Schematic of a mouse body with color-coded paws and polar plots indicating the phase of the step cycle in which each paw is positioned to reference FL paw (red). Each radial axis represents a walking speed. FL: front-left, HL: hind-left (orange), FR: front-right (blue), HR: hind-right (purple). Quantification of gain and phase of baseline compensatory eye movements: **d**, the optokinetic reflex (OKR), **e**, the vestibular-ocular reflex (VOR) and **f**, the visually enhanced VOR (VVOR) in *Pcp2^+/+^;Kir2.1* (n = 11 mice) and *Pcp2^cre/+^;Kir2.1* (n = 11 mice) groups. Mixed-effects analysis \**P* < 0.05, \*\**P* < 0.01. **g**, Quantification of VOR gain reversal training in *Pcp2^+/+^;Kir2.1* (n = 7 mice) and *Pcp2^cre/+^;Kir2.1* (n = 7 mice) groups. Mixed-effect analysis \*\*\**P* < 0.001. **h**, Conditioned responses amplitude and **i**, fraction of eyelid closure in *Pcp2^+/+^;Kir2.1* (n = 16 mice) and *Pcp2^cre/+^;Kir2.1* (n = 10 mice) groups. Mixed-effects analysis and two-way ANOVA \*\*\**P* < 0.001. **j**, Onset and peak time of the unconditioned response (UR) in *Pcp2^+/+^;Kir2.1* (n = 16 mice) and *Pcp2^cre/+^;Kir2.1* (n = 10 mice) groups. Mann-Whitney U test and Unpaired *t*-test. Data represent mean ± SEM.

**Supplementary Fig. 4:**
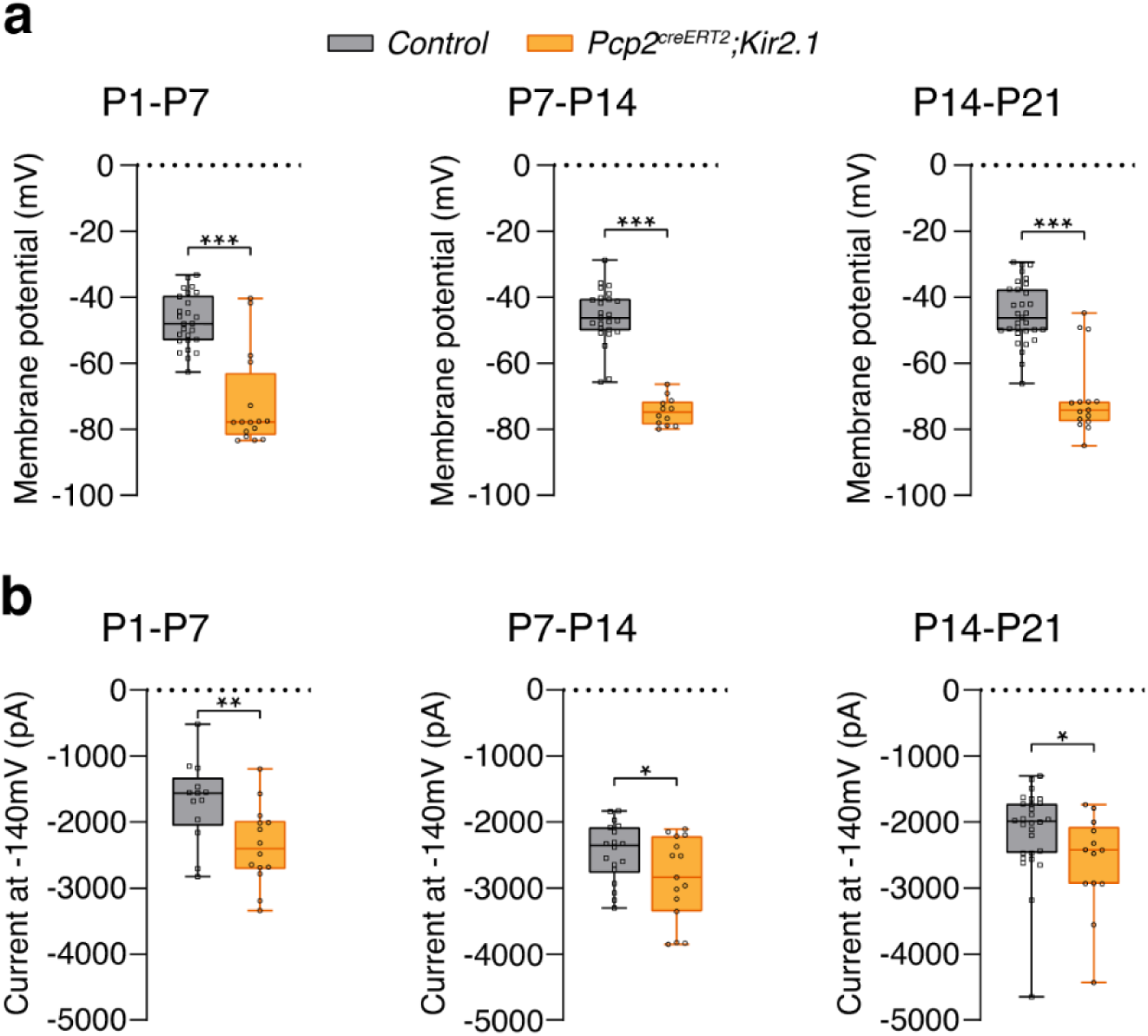
Overexpression of Kir2.1 decreases the excitability of Purkinje cells at different developmental ages. **a**, Membrane potential at P7 in control (n = 26 cells/3 mice) and *Pcp2^creERT2^;Kir2.1* (n = 16 cells/4 mice), at P14 in control (n = 25 cells/2 mice) and *Pcp2^creERT2^;Kir2.1* (n = 12 cells/5 mice) and at P21 in control (n = 13 cells/3 mice) and *Pcp2^creERT2^;Kir2.1* (n = 14 cells/5 mice) Purkinje cells. Unpaired *t*-test with Welch’s correction and Mann-Whitney *U* test \*\*\**P* < 0.001. **b**, Current amplitude hyperpolarized at −140mV at P7 in control (n = 13 cells/3 mice) and *Pcp2^creERT2^;Kir2.1* (n = 14 cells/4 mice), at P14 in control (n = 18 cells/6 mice) and *Pcp2^creERT2^;Kir2.1* (n = 15 cells/5 mice) and at P21 in control (n = 28 cells/7 mice) and *Pcp2^creERT2^;Kir2.1* (n = 13 cells/4 mice) Purkinje cells. Unpaired *t*-test and Mann-Whitney *U* test \**P* < 0.05, \*\**P* < 0.01.

**Supplementary Fig. 5:**
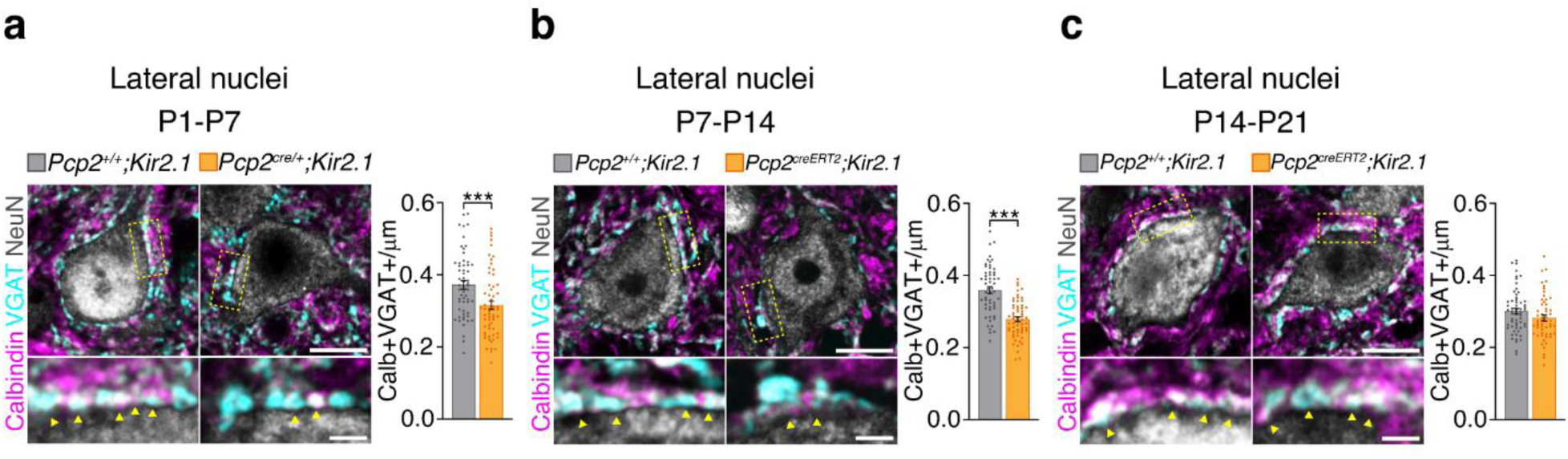
Overexpression of Kir2.1 decreases the number of inhibitory inputs Purkinje cells make on the lateral nuclei cells in the first postnatal weeks. **a**, Density of Calb+VGAT+ puncta contacting NeuN+ neurons at P7 in *Pcp2^+/+^;Kir2.1* (n = 58 cells/3 mice) and *Pcp2^cre/+^;Kir2.1* (n = 60 cells/3 mice) groups. Mann-Whitney *U* test *** *P* < 0.001. **b**, Density of Calb+VGAT+ puncta contacting NeuN+ neurons at P14 in *Pcp2^+/+^;Kir2.1* (n = 55 cells/3 mice) and *Pcp2^creERT2^;Kir2.1* (n = 59 cells/3 mice) groups. Unpaired *t*-test with Welch’s correction \*\*\**P* < 0.001. **c**, Density of Calb+VGAT+ puncta contacting NeuN+ neurons at P21 in *Pcp2^+/+^;Kir2.1* (n = 59 cells/3 mice) and *Pcp2^creERT2^;Kir2.1* (n = 55 cells/3 mice) groups. Unpaired *t*-test. Representative images (top) and insets (bottom) of Purkinje cell Calbindin+ axonal terminals (magenta) and presynaptic VGAT+ puncta (cyan) surrounding a NeuN+ neuron (grey) in the lateral nuclei of *Pcp2^+/+^;Kir2.1* and *Pcp2^cre/+^;Kir2.1* mice (**a**, **b**, **c**). Scale bar Top = 10µm, Bottom = 2 µm. Bar data represent mean ± SEM.

**Supplementary Fig. 6:**
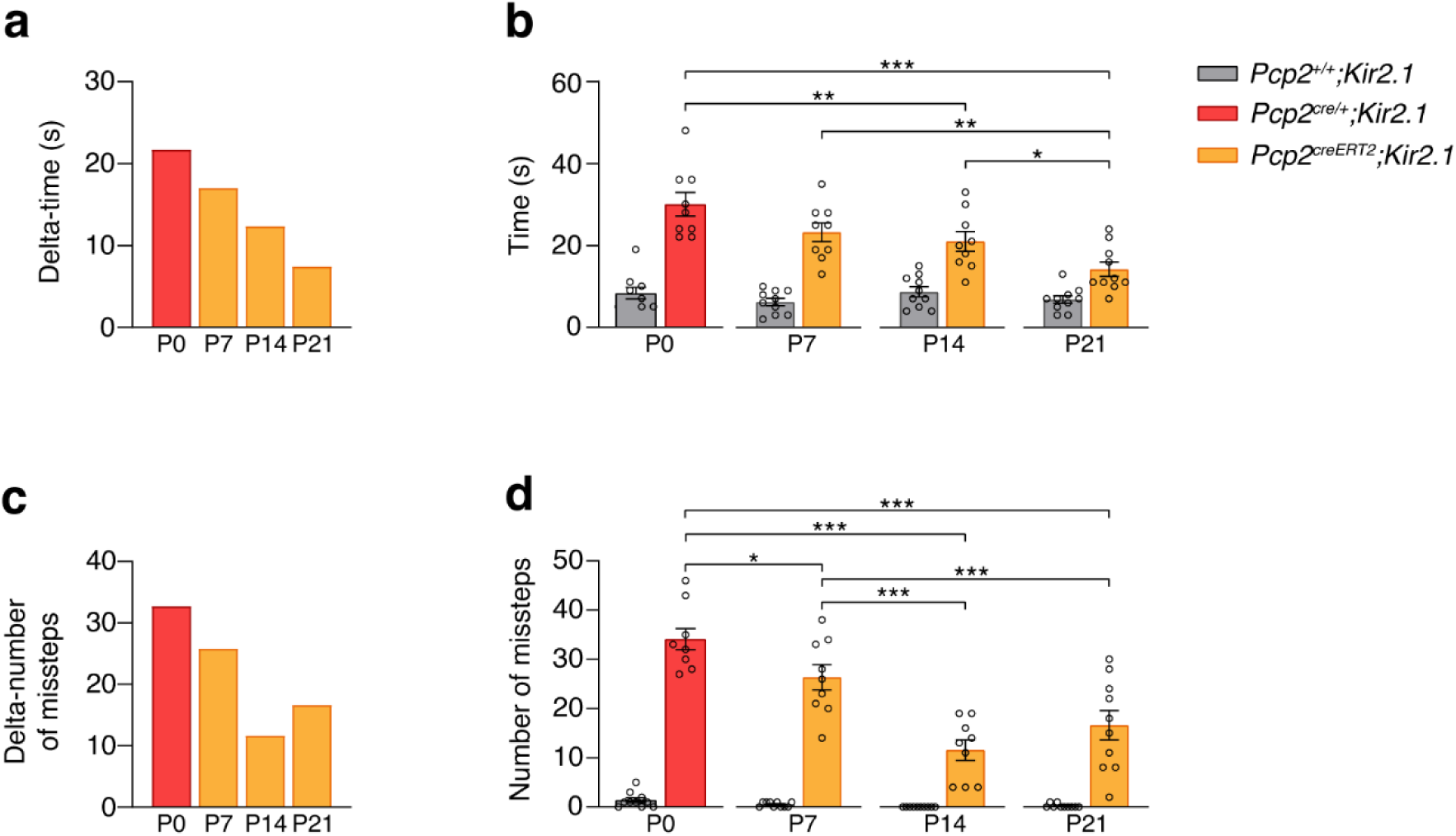
Relative effects of Purkinje cell activity reduction in balance beam performance. **a**, The time animals take to cross the beam is represented by delta-time (the difference between mutant and control time to cross the beam). **b**, The time to cross the beam in different experimental time points. Mixed-effects analysis followed by Tukey’s multiple comparisons \**P* < 0.05, \*\**P* < 0.01, *** *P* < 0.001. **c**, The number of missteps animals make while crossing the beam is represented by the delta-number of missteps (difference between mutant and control number of missteps). **d**, The number of missteps animals make while crossing the beam in different experimental time points. Mixed-effects analysis followed by Tukey’s multiple comparisons \**P* < 0.05, *** *P* < 0.001. Ages represent the starting point of Kir2.1 expression in the mutant mice.

**Supplementary Fig. 7:**
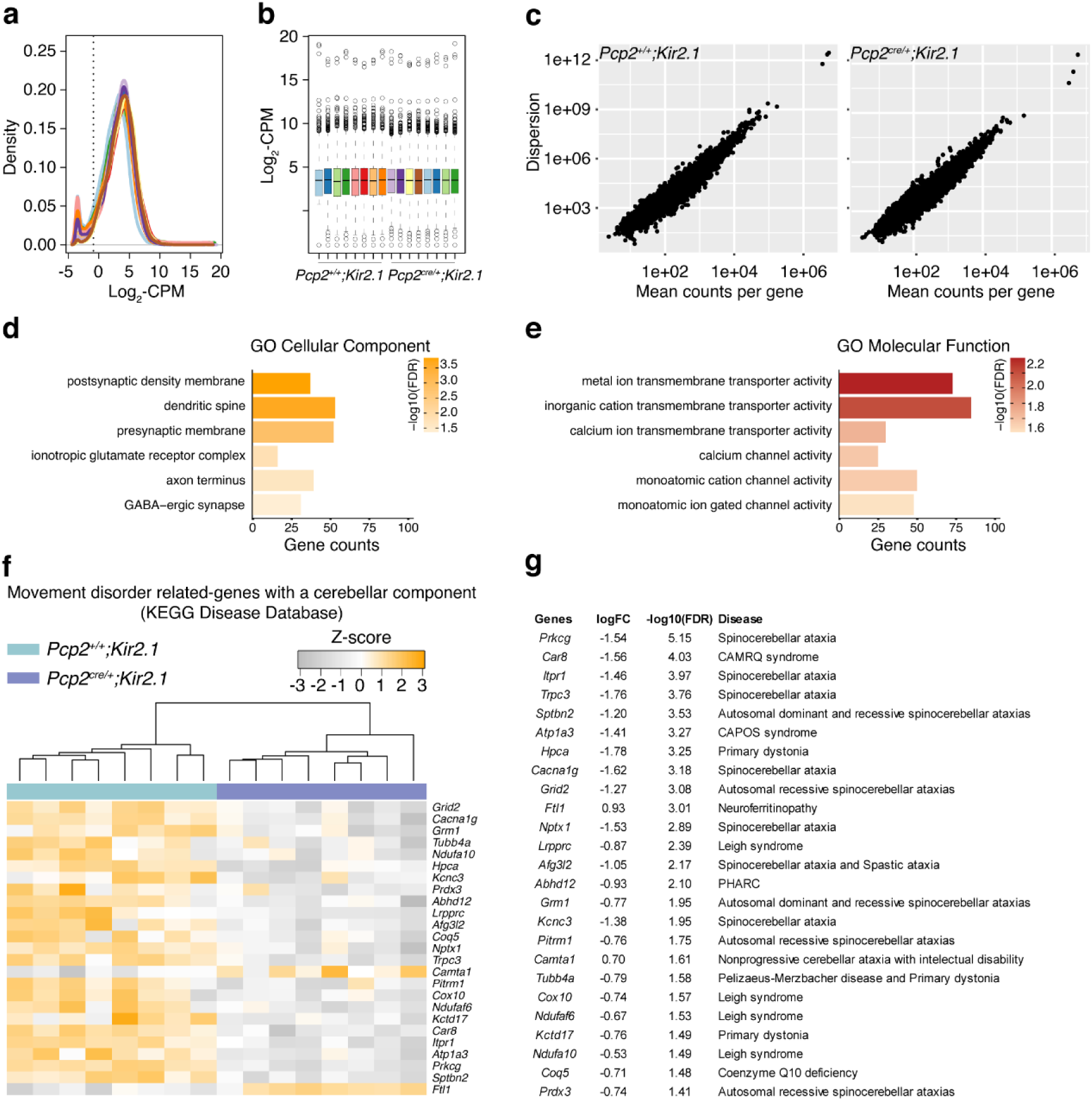
Validation of the RNA sequencing experiments, gene ontology analysis, and movement disorder related-genes. **a**, Comparison of log_2_-CPM density across samples (n = 8 mice per group). **b**, Boxplots of log_2_-CPM values showing mRNA expression distributions for each sample. **c**, Dispersion patterns of counts per gene in the different groups. Selected gene ontology (GO) enrichment analysis for the differentially expressed genes (DEG) **d**, Cellular Component, and **e**, Molecular Function categories plotted by gene count and –log_10_(FDR). **f**, Heatmap representing relative levels (Z-scores) of common genes between DEG from *Pcp2^cre/+^;Kir2.1* Purkinje cells at P7 when compared with the control, and genes related to movement disorders with a cerebellar component from the KEGG Disease Database. Samples and genes have been reordered by the method of hierarchical clustering. **g**, List of 25 common genes, log fold-change, −log10(FDR), and disorder they associate with. CPM: counts per million; FDR: false discovery rate. CAMRQ: Cerebellar ataxia-intellectual disability-disequilibrium syndrome. CAPOS: cerebellar ataxia, areflexia, pes cavus, optic atrophy, and sensorineural hearing loss syndrome. PHARC: polyneuropathy, hearing loss, ataxia, retinitis pigmentosa, and cataract.

**Supplementary Fig. 8:**
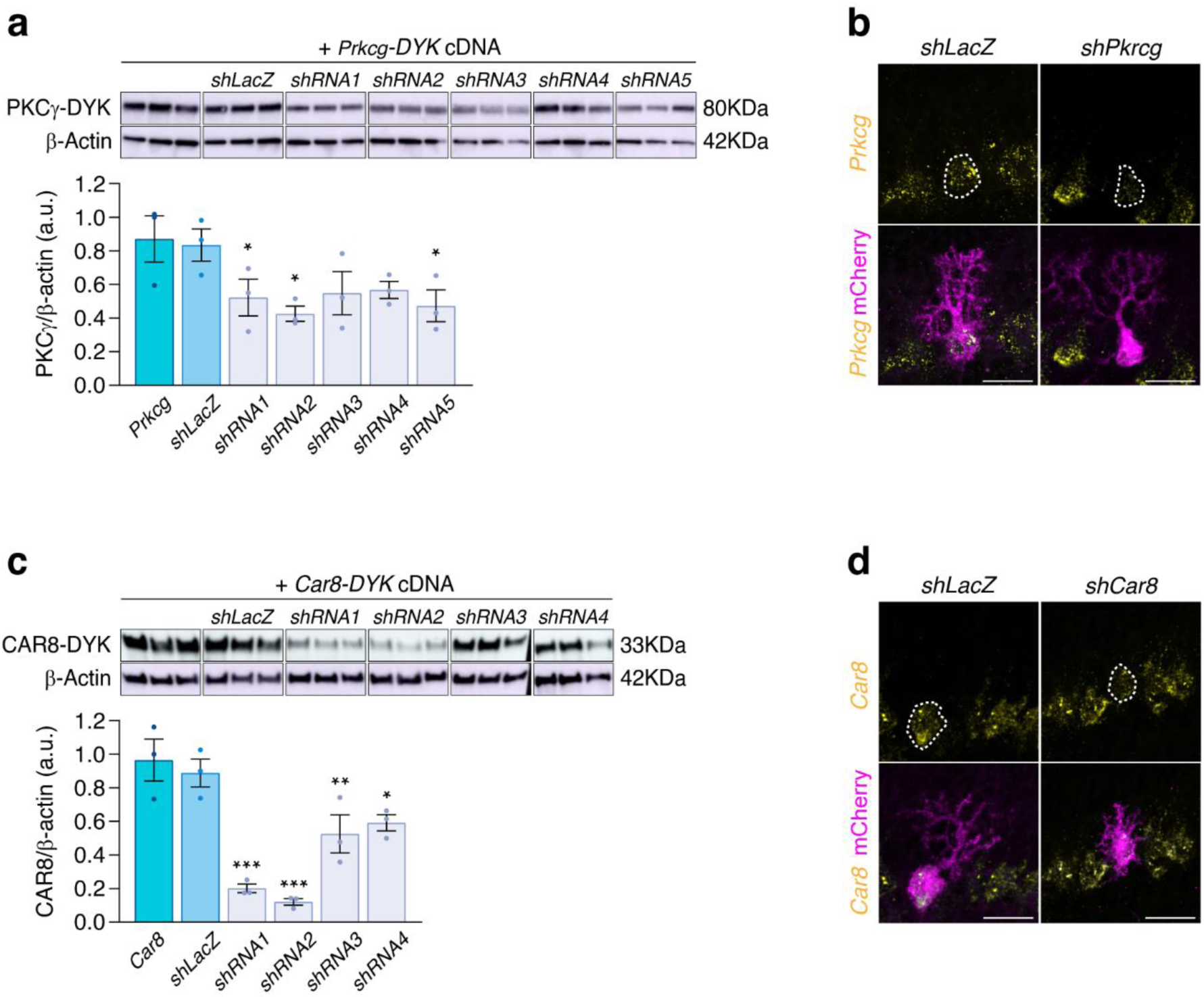
Selection of *Prkcg* and *Car8* shRNA sequences and validation of their efficacy by *in situ* hybridization. **a**, Western blot detection and quantification of DYK-tagged PKCγ normalized to β-actin from HEK cells transfected with *Prkcg-DYK* cDNA plasmid without or with *shLacZ* plasmid or one of five *shRNA* plasmids against *Prkcg* (n = 3). One-way ANOVA with multiple comparisons * *P* < 0.05. **b**, Representative images of RNAscope method showing the expression of *Prkcg* (yellow) and mCherry (magenta) in *shLacZ* and *shPrkcg* conditions from P7 *Pcp2^cre/+^* Purkinje cells. **c**, Western blot detection and quantification of DYK-tagged CAR8 normalized to β-actin from HEK cells transfected with *Car8-DYK* cDNA plasmid without or with *shLacZ* plasmid or one of four *shRNA* plasmids against *Car8* (n = 3). One-way ANOVA with multiple comparisons: * *P* < 0.05, \*\**P* < 0.01, *** *P* < 0.001. **d**, Representative images of RNAscope method showing the expression of *Car8* (yellow) and mCherry (magenta) in *shLacZ* and *shCar8* conditions from P7 *Pcp2^cre/+^* Purkinje cells. Scale bar = 25 µm (**b**, **d**). Data represent mean ± SEM.

Supplementary video 1. Balance beam test in *Pcp2^+/+^;Kir2.1* adult mouse.

Supplementary video 2. Balance beam test in *Pcp2^cre/+^;Kir2.1* adult mouse

Supplementary video 3. Accelerated rotarod test in *Pcp2^+/+^;Kir2.1* and *Pcp2^cre/+^;Kir2.1* adult mice.

Supplementary video 4. LocoMouse test in *Pcp2^+/+^;Kir2.1* adult mouse.

Supplementary video 5. LocoMouse test in *Pcp2^cre/+^;Kir2.1* adult mouse.

## Methods

### Mouse strains

The following transgenic mouse lines were used in this study and maintained in a C57BL/6 background (Charles River Laboratories): *Pcp2^creERT2^* (*Tg(Pcp2-creER^T2^)17.8.ICS*)^22^, *Ai14* (*B6;129S6-Gt(ROSA*)26Sor_tm14(CAG-tdTomato)Hze/J)_24, *Kir2.1 (Gt(ROSA)26Sor_tm2(CAG-KCNJ2/mCherry)Fmr)_*_23_ and *Pcp2_cre/+_(B6.Cg-Tg(Pcp2-cre)3555Jdhu/J)*^77^. The *Ai14* (Jackson Laboratories #007908) and *Kir2.1* (kindly provided by Guillermina López-Bendito) mice were crossed with inducible *Pcp2^creERT2^* to drive expression of *tdTomato* or *Kir2.1-mCherry* in Purkinje cells, respectively, upon injection of tamoxifen. Tamoxifen (Sigma-Aldrich, Cat. #T5648, 20 mg/ml in corn oil) induction of Cre recombinase was administered via intraperitoneal injection into postnatal day (P)1, P7, P14 or P21 *Pcp2^creERT2^*;*Ai14* or *Pcp2^creERT2^; Kir2.1* pups. For space recombination of Purkinje cells, 100mg/Kg of body weight of tamoxifen was injected in P1 pups, 5mg/Kg in P7 pups, and 1mg/Kg in P14 pups. For full recombination of Purkinje cells, 100mg/Kg of tamoxifen for three consecutive days was injected at P7, P14, or P21. The *Kir2.1* mouse line was crossed with *Pcp2^cre/+^* to conditionally express the Kir2.1 channel in all Purkinje cells. For *in utero* electroporation surgeries, FVB/NHsd (Envigo) time-mated female mice were used and housed individually. The day of the vaginal plug was stipulated as embryonic day (E)0.5. Both male and female mice were used in all experiments. All animals were maintained under standard, temperature-controlled laboratory conditions. Mice were kept on a 12:12 light/dark cycle and received water and food *ad libitum*. All procedures were approved by the Dutch Ethical Committee for Animal Experiments and conducted in accordance with the Institutional Animal Care and Use Committee of Erasmus Medical Center (IACUC Erasmus MC), the European and the Dutch National Legislation.

### *in utero* electroporation

Embryonic (E)12.5 timed-pregnant FVB/NHsd females were deeply anesthetized with isoflurane (5% induction, 2% maintenance) in oxygen at a flow rate of 1 l/min and the body temperature was maintained at 37 °C. Buprenorphine (0.1mg/kg) was administered for analgesia via subcutaneous injection; the abdominal cavity was cut open, and the uterus was exposed, making the embryos accessible. A beveled glass micropipette was used to inject a combination of 3μg/μl of *pCAG-GFP* (Addgene; #11150; a gift from C. Cepko^78^) and 1μg/μl of *pCAG-Kir2.1-T2A-tdTomato* (Addgene; #60598; a gift from M. Scanziani^79^) with 0.05% of FastGreen in the fourth ventricle of mouse embryos. Five pulses of current (35 V for 50 ms, spaced 900 ms) were delivered via a 2mm tweezertrode (CUY650P2, Nepa Gene) connected to an electroporator (BTX, ECM830). The electrodes were positioned to target Purkinje cell progenitors in the cerebellar ventricular zone. After electroporation, the uterus was placed back in the abdominal cavity, the abdominal wall and skin were sutured and the female was allowed to recover.

### Immunohistochemistry

Mice were deeply anesthetized with sodium pentobarbital by intraperitoneal injection, transcardially perfused with sodium chloride solution followed by 4% paraformaldehyde (PFA) in 0.1M phosphate buffer (PB) (pH 7.4). Dissected brains were then fixed for 2h at 4°C in the same solution and cryoprotected in 30% sucrose/PB overnight at 4°C. The next day, the brains were embedded in 14% gelatin/ 30% sucrose/PB solution, fixed for 2h at room temperature, and incubated overnight at 4°C in 30% sucrose/0.1M PB solution. A freezing microtome was used to cut 50µm sagittal sections. Free-floating sections were rinsed with PB and processed for antigen retrieval by being incubated for 2h in a solution of 10mM of sodium citrate at 80°C. After, sections were rinsed and incubated in a blocking solution containing 0.5% Triton X-100 and 10% normal horse serum in PB for 2h at room temperature to block nonspecific protein-binding sites. Subsequently, sections were incubated with primary antibodies in antibody solution (0.5% Triton X-100 and 2% normal horse serum in PB) overnight at 4°C. The following primary antibodies were used in this study: rabbit anti-RFP (1:1000, Rockland, #600-401-379), chicken anti-RFP (1:1000, Rockland, #600-901-379), goat anti-GFP (1:1000, Rockland, #600-101-215), mouse anti-calbindin D-28K (1:10000, Swant, #300), guinea pig-VGAT (1:500, Synaptic Systems, #131004) and rabbit-NeuN (1:1000, Millipore, #ABN78).

Sections were then rinsed in PB and incubated with secondary antibodies for 2h at room temperature. Secondary antibodies used in this study were: Cy3-AffiniPure Donkey anti-Rabbit (1:1000, #711-165-152), Cy3-AffiniPure Donkey anti-Chicken (1:1000, #703-165-155), Alexa Fluor 488-AffiniPure Donkey anti-Goat (1:1000, #705-545-147), Alexa Fluor 488-AffiniPure Donkey anti-Mouse (1:1000, #715-545-150), Cy3-AffiniPure Donkey anti-Mouse (1:1000, #715-165-150), Alexa Fluor 488-AffiniPure Donkey anti-Guinea Pig (1:1000, #706-545-148) and Cy5-AffiniPure Donkey anti-Rabbit (1:1000, #711-175-152) all from Jackson ImmunoResearch. Finally, brain sections were counterstained with the fluorescent nuclear dye 4′,6-diamidino-2-phenylindole (DAPI) (Thermo Scientific, #D3571) and mounted with Mowiol (Polysciences, Inc, #17951).

### Single-molecule fluorescence *in situ* hybridization

Mice were perfused as described above, and brains were postfixed for 2h at 4°C and cryoprotected in 30% sucrose/PB overnight at 4°C. Sagittal sections of 30µm were obtained using a freezing microtome, mounted in RNAse-free SuperFrost Plus slides (Epredia), and probed against the candidate genes and mCherry according to the manufacturer protocol. RNAscope Multiplex Fluorescent Reagent Kit v2 (323100), probes *Mm-Car8-C2* (#514171), *Mm-Prkcg-C2* (#417911) from ACDBio were used together with rabbit anti-RFP (1:1000, Rockland, #600-401-379) and Cy3-AffiniPure Donkey anti-Rabbit (1:1000, Jackson ImmunoResearch, #711-165-152).

### Image acquisition and analysis

Images were acquired with a 1024x1024 pixel resolution and 8 bit depth with an LSM 700 (Carl Zeiss Microscopy) confocal laser scanning microscope or an Axio Imager.M2 (Carl Zeiss Microscopy, LLC, USA). For each experiment, images were acquired using the same laser power, detection filter settings, pinhole, and photomultiplier gain. To image individual Purkinje cells, sagittal sections images were acquired with 20X/0.8NA or 40X/1.3NA oil objective with a z-interval of 1µm. Due to the morphology variability in different cerebellar regions, individual Purkinje cells were imaged across the cerebellar cortex. To quantify the complexity of dendrite arborization in individual Purkinje cells, the maximum projection of z-stack images of each cell was analyzed with the Sholl analysis macro implemented in FIJI (ImageJ) software^80^. The longest dendrite length was obtained as the result of Sholl analysis. To quantify the Purkinje cell area, the maximum projection of each image was thresholded in FIJI (ImageJ) to fit the area of the cell and measured. The molecular layer (ML) ratio was calculated as the percentage of Purkinje cell height (the distance from the edge of the Purkinje cell soma to the apical edge of the molecular layer) per the molecular layer height. The number of neurites was measured as the number of processes coming out of the soma of a Purkinje cell. To image presynaptic puncta, coronal sections images were acquired with 63X/1.4NA oil objective with a z-interval of 1µm. To quantify presynaptic densities, the perimeter of a NeuN-positive soma from cerebellar nuclei cells was measured using FIJI (ImageJ). Presynaptic puncta Calbindin-positive and VGAT-positive were detected automatically using the “Find maxima” tool to count the number of puncta colocalizing with soma perimeter. The number of Calbindin-positive colocalizing with VGAT-positive puncta was then normalized to the perimeter (μm) of the cell analyzed.

### *ex vivo* Purkinje cell recordings and analysis

*ex vivo* slice recordings of Purkinje cells were made in cerebellar slices from a total of 52 mice at three developmental ages: P7, P14, and P21. Each age is given ± 1 day. Cerebellar tissue was collected from either *Pcp2^creERT2^;Ai14* or *Pcp2^creERT2^;Kir2.1* mice, previously injected with tamoxifen at P1, P7 or P14 to drive sparse expression of *tdTomato* reporter gene or *Kir2.1-mCherry*, respectively, in Purkinje cells. As previously described^21^, mice were deeply anesthetized with isoflurane and decapitated. The brain was rapidly dissected and placed in an ice-cold slice solution (continuously equilibrated by perfusing 95% O_2_ and 5% CO_2_) containing the following (in mM): 240 Sucrose, 2.5 KCl, 1.25 NaH_2_PO_4_, 2 MgSO_4_, 1 CaCl_2_, 26 NaHCO_3_, 10 D-glucose. Sagittal slices of vermal cerebellar tissue were cut 250µm thick in an ice-cold slicing solution using a vibratome (VT1000S, Leica Biosystems, Wetzlar, Germany) with a ceramic blade (Campden Instruments Ltd, Manchester, United Kingdom). Slices were immediately incubated in equilibrated artificial cerebrospinal fluid (aCSF) and maintained at 34°C for one hour before being transferred to room temperature. The aCSF consisted of (in mM): 124 NaCl, 5 KCl, 1.25 Na_2_HPO_4_, 2 MgSO_4_, 2 CaCl_2_, 26 NaHCO_3_, 20 D-glucose. Individual slices were transferred to a recording chamber and maintained at 34°C with a feedback temperature controller (Scientifica, Uckfield, United Kingdom) under continuous perfusion with equilibrated aCSF. For some recordings, the aCSF was supplemented with 300 µM barium (Sigma-Aldrich, Merck KGaA, Darmstadt, Germany). Purkinje cells were selected based on their expression of tdTomato for *Pcp2^creERT2^;Ai14*, no expression of mCherry for *Pcp2^creERT2^;Kir2.1,Ctl* and expression of mCherry for *Pcp2^creERT2^;Kir2.1*. Neurons were visualized with SliceScope Pro 3000, a CCD camera, a trinocular eyepiece (Scientifica, Uckfield, UK), and an ocular (Teledyne Qimaging, Surrey, Canada). Whole-cell recordings were obtained using borosilicate pipettes (Harvard apparatus, Holliston, MA, USA) with a resistance of 4-6 MΩ, filled with internal solution consisting of (in mM): 9 KCl, 3.48 MgCl_2_, 4 NaCl, 120 K^+^-Gluconate, 10 HEPES, 28.5 Sucrose, 4 Na_2_ATP, 0.4 Na_3_GTP in total pH 7.25-7.35, osmolarity 290-300 mOsmol/Kg (Sigma-Aldrich, Merck KGaA, Darmstadt, Germany). The internal solution was supplemented with 1 mg/ml biocytin to allow for histological examination of recorded Purkinje cells. Recordings were acquired with Patchmaster (HEKA Electronics, Lambrecht, Germany) using an ECP-10 amplifier (HEKA Electronics, Lambrecht, Germany) and digitized at 20 kHz. Data files were imported into and analyzed in MATLAB (Mathworks, Natick, MA, USA). Before the whole-cell recording configuration, the capacitance was optimally compensated. Whole-cell recordings were performed in Purkinje cells that were held at −65 mV. The input resistance was assessed by a −10 mV test pulse. Recordings with a series resistance greater than 25 MΩ were excluded. After achieving whole-cell configuration, zero current was injected in the current-clamp configuration to assess the presence or absence of spontaneous action potential firing and the resting membrane potential, defined as the mode of the resulting trace. The current-voltage (I-V) curve was generated in voltage-clamp mode by injecting increasing 500-ms voltage steps from −140 mV to 0 mV at 10-mV increments and measuring the mode of the resulting current.

### Behavioral tests

All the behavioral tests were performed in adult mice (P60 to P90).

#### Balance beam test

Mice were tested on a 1-meter-long 12mm width beam with a flat surface. The beam was located between two elevated platforms 50 cm above the surface. A cage was placed at the end of the beam as a finish point. The test consisted of two training days and one testing day. Mice were trained for three trials per day. Any trial in which a mouse fell 5 consecutive times was considered completed. Motor coordination was determined by averaging the time taken to cross the beam and the number of missteps per run^81^.

#### Accelerating rotarod test

Mice were tested in an accelerating rotarod in 4 non-consecutive trials during 5 consecutive days. From days 1 to 4, the rotating rod was accelerated to 40 rpm, while on the 5th day, the maximum speed was increased to 80 rpm. Mice were given an hour to rest between trials. A trial was considered completed when the mouse reached a maximum time of 300 s on the rod, fell off the rod, or rotated 3 consecutive times. Motor coordination was determined by averaging the latency trials to fall on the accelerating rod (Ugo Basile Biological Research Apparatus).

#### LocoMouse

The LocoMouse test was performed to assess whole body coordination during overground locomotion^29^. Briefly, mice crossed a glass corridor (66.5 cm long, 4.5 cm wide, and 20 cm high) between two dark boxes. Below the corridor, a mirror (66 cm × 16 cm) was placed at 45° to allow for simultaneous collection of side and bottom views. Side and bottom views of walking mice were recorded with a high-speed camera (Basler, 1440 x 250 pixels, 400 fps). The test consisted of two consecutive habituation days followed by three consecutive experimental days. On the habituation days, mice walked freely between the two dark boxes for 15 min. On each experimental day, 15 trials per animal were collected. Individual trials consisted of single crossings of the corridor, and animals were not required to walk continuously throughout a trial. A DeepLabCut network was trained using the side camera view of the LocoMouse to track the paws, nose and tail base of each individual mice^82^. Data was then processed using a custom-written Python (v3.7) code^30^.

#### Open field test

The open field test was performed in a white-opaque polypropylene squared box (50 x 50 x 40 cm) under uniform lighting conditions. Each animal was placed in the center of the arena and allowed to explore for 10 min. A digital video system recorded the overall activity inside the box (Carl Zeiss Tessar 2.0/3.7 2MP V-UBM46 Autofocus Camera). Data was analyzed using the OptiMouse, an open-source MATLAB program^83^The software automatically calculated each mouse’s total distance traveled and speed to evaluate spontaneous locomotor activity.

#### Compensatory eye movements

Compensatory eye movement recordings were performed as previously described^21^. Briefly, mice were surgically prepared for head-restrained recordings by placing a custom-built metal pedestal on their frontal and parietal bones under general anesthesia with isoflurane/O_2_. Three days after recovering from the surgery, mice were head-fixed and placed in a mouse holder in the center of a turntable (60 cm diameter), surrounded by a cylindrical screen (63 cm diameter) with a random-dotted pattern (*drum*). Eye movements were recorded with a CCD camera fixed to the turntable, and eye-tracking software was used (ETL-200, ISCAN systems, Burlington, NA, USA). Eyes were illuminated using two table-fixed infrared emitters and a third emitter fixed to the camera, which produced a tracked corneal reflection used as the reference point. The recording camera was calibrated by being moved left-right by 20° peak to peak during periods that the eye did not move^84^. Gain and phase values of eye movements were calculated using custom-made MATLAB scripts, available at GitHub (GitHub, San Francisco, CA, USA, https://github.com/MSchonewille/iMove;^85^). Mice were familiarized with the experimental set-up three days before the experiments by providing visual and vestibular stimulations for 30 min. Compensatory eye movements were induced using a sinusoidal rotation of the drum in light (optokinetic reflex-OKR), rotation of the table in the dark (vestibular ocular reflex-VOR), or the rotation of the table in the light (visual VOR-VVOR) with an amplitude of 5° at 0.1–1 Hz. To assess motor learning, mice were subjected to a mismatch between visual and vestibular input to adapt the VOR. The ability to perform VOR phase-reversal was tested during a period of 5 days, consisting of six 5-minute training sessions every day with VOR recordings before, between, and after the training sessions. Mice were kept in the dark between recording sessions to avoid unlearning the adapted responses. On the first training day, the visual (the drum) and vestibular stimuli (turntable) rotated in phase at 0.6 Hz and both with an amplitude of 5°, inducing a decrease of gain. In the following days, the drum amplitude was increased to 7.5° (day 2) and 10° (days 3, 4, and 5), while the amplitude of the turntable remained at 5°. This resulted in the reversal of the VOR direction, an inversion of the compensatory eye movement driven by vestibular input, moving the eye in the same direction as the head rotation instead of the normal compensatory opposite direction. Motor performance in response to the different stimulations was measured by calculating the response’s gain (ratio of eye movement amplitude to stimulus amplitude) and phase (eye to stimulus difference in degrees).

#### Eyeblink conditioning

Eyeblink conditioning was performed as previously described^86^. Briefly, a small metal pedestal with a square magnet on top was placed on the mice skull under general anesthesia with 2% isoflurane/O_2_. Three days after recovering from the surgery, mice were head-fixed and placed on top of a foam cylindrical treadmill on which they could walk freely in custom-built sound- and light-attenuating boxes. Eyelid movements were monitored under infrared illumination using a high-speed (333 frames/s) monochrome Basler video camera (Basler ace 750-30gm). All stimuli and measurement devices were controlled by National Instrument hardware and custom-written LabVIEW software. The conditioned stimulus (CS) was a blue LED light (CS duration 280 ms, LED diameter 5 mm) placed 10 cm in front of the mouse’s head. The unconditioned stimulus (US) consisted of a 30 ms duration mild corneal air puff, which was controlled by a VHS P/P solenoid valve (Lohm rate, 4750 Lohms; Internal volume, 30µl, The Lee Company®, Westbrook, USA) and delivered via a 27.5mm gauge needle that was perpendicularly positioned at about 5 mm from the center of the left cornea. The back pressure on the solenoid valve was set at 30 psi. Mice were habituated for one day to the set-up (two 30 min sessions spanning 6 hours) prior to the training. No stimuli were delivered during habituation. To assess motor learning, mice were subjected to eyeblink conditioning training for 5 days, two sessions per day with a resting period of 6 hours. Before the first training session, each animal underwent one baseline session to establish that the CS did not elicit any reflexive eyelid closure. This baseline session consisted of 15 CS-only trials and 3 US-only trials. Immediately after this, the training sessions begun. During each session, every animal received a total of 200 paired CS-US trials, 20 US-only trials, and 20 CS-only trials. These trials were presented over 20 blocks, and each block consisted of 1 US-only trial, 10 paired CS-US trials, and 1 CS-only trial. The interval between the onset of the CS and that of the US was set at 250 ms. All experiments were performed at approximately the same time of the day by the same experimenter. Individual eyeblink traces were analyzed with a custom-written MATLAB script (R2018a, Mathworks). First, the 2000 ms eyeblink traces were imported from the MySQL database into MATLAB. The trials were aligned at zero for the 500 ms pre-CS baselines. Trials with significant activity in the 500 ms pre-CS period (> 7 times the interquartile range) were considered invalid and disregarded for further analysis. The eyelid signal was min-max normalized so that a fully open eye corresponded with a value of 0 and a fully closed eye with a value of 1^87^. In valid normalized CS-only trials, eyelid responses were considered as a conditioned response (CR) if the maximum amplitude was larger than 0.10 in the interval between 100-500 ms after CS onset and the presence of a positive slope in the 150 ms before the time point where the US would have been delivered (US is omitted in CS-only trials)^88^.

### RNA sequencing and differential expression analysis

#### Purkinje cell collection

To isolate Purkinje cells, the cerebellar tissue was extracted from either P7 *Pcp2^+/+^;Kir2.1* or P7 *Pcp2^cre/+^;Kir2.1* mice. Sagittal slices of vermal cerebellar tissue were obtained and maintained as described in the *ex vivo* Purkinje cell recordings section. For cell collection, individual slices were transferred into a recording chamber and maintained at 34°C under continuous perfusion with equilibrated aCSF. For P7 *Pcp2^cre/+^;Kir2.1* mice, Purkinje cells were selected based on their expression of Kir2.1-mCherry. Neurons were extracted into a glass pipette using negative pressure and 50 cells were sampled across the cerebellar cortex. For each genotype, eight biological samples were used for RNA sequencing. Cells were collected in a mild hypotonic lysis buffer composed of 0.2% Triton-X and 2U/µl of RNAse inhibitor. Isolated cells were kept on ice during the collection process or immediately stored at −80°C for RNA extraction.

#### cDNA library preparation and RNA sequencing

Library preparation and RNA sequencing experiments were performed by the Erasmus Center for Biomics (Erasmus MC, Rotterdam, The Netherlands). For each biological sample, cDNA libraries were generated using the Smart-seq2 method^89^. In brief, cells were lysed, and polyA+ RNA was reverse-transcribed using an oligo(dT) primer. Template switching by reverse transcriptase was achieved by using an LNA containing TSO oligo. The reverse transcribed cDNA was pre-amplified with primers for 18 cycles, followed by clean-up. Tagmentation was performed on 1 ng of the pre-amplified cDNA using Illumina Nextera DNA Flex kit. The tagmented library was extended with Illumina adaptor sequences by PCR for 12 cycles and purified. The resulting sequencing library was measured on Bioanalyzer and equimolar loaded onto a flowcell and sequenced according to the Illumina TruSeq v3 protocol on the Illumina HiSeq2500 platform with a single read 50 base-pairs and dual 9 base-pairs indices.

#### Data processing

For data analysis, Illumina adapter sequences and poly-A stretches were trimmed from the reads. The remaining sequences were mapped to the mouse reference sequence GRCm38 (mm10) using HISAT2 (version 2.1.0). From the alignments, the reads per gene were determined with the htseq-count program (version 0.11.2). Gene and transcript information was obtained from Ensembl (build 101) using ccds genes. Count data were then analyzed using R studio version 0.99.484 according to the edgeR-limma package workflow^90^. Briefly, Ensembl IDs were annotated to Entrez gene IDs, gene symbols, gene names, and chromosomes using the Mus.musculus package. Next, raw counts were transformed into counts per million (CPM) and log_2_-CPM, and genes with a CPM below 0.2 in at least three samples were filtered out. Normalization of gene expression distributions was performed using the Trimmed Mean of M-values method.

#### Analysis of differentially expressed genes (DEG)

Differentially expressed genes between Purkinje cells from *Pcp2^+/+^;Kir2.1* and *Pcp2^cre/+^; Kir2.1* were plotted by the method of hierarchical clustering heatmap using normalized row z-scores in R. To identify the most biologically significant genes, data were organized in a volcano plot with a fold change cut off of 0.5 and a –log_10_(FDR) cut off of 1.3. Gene ontology (GO) analysis were carried out using the Gene Ontology tool (http://pantherdb.org/) and plotted in R. DEGs were selected for GO analyses. The total list of genes identified in the biological samples was used as the reference list. GO terms with corrected *P*<0.05 were considered significantly enriched

#### Selection of candidate genes

A list with 216 related human movement disorders genes was extracted from the KEGG Disease database (https://www.genome.jp/kegg/disease/). This database collects known genetic and environmental factors of diseases. We compared this list to the 1602 DEG from the RNA sequencing data obtained from P7 *Pcp2^+/+^;Kir2.1* vs. P7 Pcp2cre/+;Kir2.1 Purkinje cells and identified 25 common genes. These were organized by −log_10_(FDR) and log fold change. The top hits were *Prkcg* and *Car8* genes.

### Cell Culture and Transfection

HEK293T cells were cultured in Dulbecco’s Modified Eagle’s medium supplemented with high glucose, Glutamax and pyruvate (31966-047, ThermoFisher Scientific). In addition, 10% fetal bovine serum, 1% of 10,000 units/ml of penicillin, and 10,000 µg/ml of streptomycin. The cultures were maintained at 37°C in a humidified atmosphere containing 5% CO_2_. HEK293T cells were plated in 6-well plates at a density of 0.5×10^6^ cells/well. On the following day, cells were transfected with plasmids expressing the full-length genes of interest and respective shRNA expressing plasmids with a 1:3 ratio using Lipofectamine 2000 (11668019, ThermoFisher Scientific). The plasmids used to expressed *Prkcg* and *Car8* were *pcDNA3.1+/C- (K)DYK-Prkcg* (OMu17867, GenScript) and *pcDNA3.1+/C-(K)DYK-Car8* (OMu00430, GenScript). shRNA sequences targeting *Prkcg* and *Car8* were designed by The RNAi Consortium (TRC) and acquired by Horizon Discovery. The antisense sequence and respective clone IDs are as follows: for *Prkcg shRNA1* 5’ TTG ATG GCA TAG AGT TCG TCG 3’ (TRCN0000022684), *shRNA2* 5’ TAA TAT GGA TCT CAT CCG ACG 3’ (TRCN0000022685), *shRNA3* 5’ ATA GGC AAT GAT CTC AGG TGC 3’ (TRCN0000022686), *shRNA4* 5’ TTC ACA TAA GTA AAG CCC TGG 3’ (TRCN0000022687) and *shRNA5* 5’ AAA CGA GCG GTG AAC TTG TGG 3’ (TRCN0000022688). For *Car8 shRNA1* 5’ AAT ATC CAG GTA ACT CCT TCG 3’ (TRCN0000114512), *shRNA2* 5’ TTT AGG TTG ATA GGT GAC TGG 3’ (TRCN0000114513), *shRNA3* 5’ TAG CAT CAG GAA ACA CTA AGC 3’ (TRCN0000114514) and *shRNA4* 5’ ATC TGG GAT ATA GTT AAA GGG 3’ (TRCN0000114515). The control *shRNA lacZ* sequence was designed as follows: 5’ AAA TCG CTG ATT TGT GTA GTC 3’.

### Western blotting

HEK293T cells were collected 24h after transfection in lysis buffer containing 50 mM Tris-HCl pH 8, 150 mM NaCl, 1% Triton X-100, 0.5% sodium deoxycholate, 0.1% SDS, and protease inhibitor cocktail. Protein concentrations were measured using Pierce BCA protein assay kit (Thermo Fisher). Samples were denatured, proteins were separated by SDS-PAGE in Criterion TGX Stain-Free Gels (Bio-Rad) and transferred onto nitrocellulose membranes with the Trans-Blot Turbo Blotting System (Bio-Rad). Membranes were blocked with 5% BSA (Sigma-Aldrich) in Tris-buffered saline (TBS)-Tween (20 mM Tris-HCl pH7.5, 150 mM NaCl and 0.1%, Tween20) for 1h and probed with the following primary antibodies: rabbit anti-DYKDDDDK-tag (1:2000, GenScript, #A00170) and mouse anti-actin (1:1000, Millipore, #MAB1501). Secondary antibodies used were goat Anti-Rabbit Immunoglobulins/HRP (1:10000, Agilent Dako, #P0448) or goat Anti-Mouse Immunoglobulins/HRP (1:10000, Agilent Dako, #P0447). Proteins were detected by the luminol-based enhanced chemiluminescence method (SuperSignal™ West Femto Maximum Sensitivity Substrate or SuperSignal™ West Dura Extended Duration Substrate, Thermo Fisher). Membranes were stripped with Restore™ PLUS Western Blot Stripping Buffer, Thermo Fisher). Densitometry of protein bands of interest was normalized to that of actin using the Image Studio Lite software (LI-COR Biosciences).

### Generation of DNA constructs

DNA vectors were generated by standard molecular biology procedures. To generate viral vectors expressing *shRNA* for the target genes, pairs of oligos were hybridized for *shlacZ* (strand1 5’ cta ggA AAT CGC TGA TTT GTG TAG TCT GAT ATG TGC AGA CTA CAC AAA TCA GCG ATT TTT TTTg 3’and strand2 5’ aat tca aaaa AAA TCG CTG ATT TGT GTA GTC TGC ACA TAT CAG ACT ACA CAA ATC AGC GAT TTc 3’), *shPrkcg* (strand1 5’ cta ggT AAT ATG GAT CTC ATC CGA CGC TCG AGC GTC GGA TGA GAT CCA TAT TAT TTTTg 3’and strand2 5’ aat tca aaaa TAA TAT GGA TCT CAT CCG ACG CTC GAG CGT CGG ATG AGA TCC ATA TTAc 3’) and *shCar8* (strand1 5’ cta ggT TTA GGT TGA TAG GTG ACT GGC TCG AGC CAG TCA CCT ATC AAC CTA AAT TTTTg 3’and strand2 5’ aat tca aaaa TTT AGG TTG ATA GGT GAC TGG CTC GAG CCA GTC ACC TAT CAA CCT AAAc 3’). Each double strand was subsequently cloned using EcoRI and AvrI sites in the *pDIO-DSE-mCherry-PSE-MCS* vector (Addgene; #129669; a gift from B. Rico^41^).

### Generation of viral particles

Viral prep AAV-FLEX-GFP was commercially acquired (Addgene; #28304-PHPeB; a gift from E. Boyden). All the other viral preps were produced as described previously^91^. In short, HEK293T cells cultured in DMEM (with 10% fetal calf serum and penicillin/streptomycin) were co-transfected with AAV2/9 serotype helper plasmid, helper plasmid pAdΔF6 and pAAV-shRNA using PEI (polyethylenimine 25kDA, linear). Seventy-two hours after transfection, the cells were harvested, lysed, and centrifuged to remove cellular debris. The AAV particles were purified from the supernatant using an iodixanol density gradient and subsequently concentrated in 5%sucrose/PBS using an Amicon Ultra-15 centrifugal filter. Viral titers were determined by quantitative PCR for the reporter mCherry present in the viral genomes (vg). mCherry primers: 5′-tcc cac aac gag gac tac ac-3′ and 5′-ctt gta cag ctc gtc cat gc-3’. The viral batches’ titers ranged from 5×10^12^ to 2×10^13^ vg/ml.

### Intraventricular viral injections in neonates

For intraventricular injections, *Pcp2^cre/+^* at P0 were cryo-anesthetized at 0°C for 1 min before the injection. Pups were placed in a custom-made plate, and a solution of AAVs diluted 1:100 with NaCl containing 0.05% of FastGreen was injected bilaterally into the lateral ventricles using a Nanoject III (Drummond Scientific Company). The injection site was placed at approximately two-fifths of the distance between the lambda and each eye, and 1μl of the viral solution was injected into each lateral ventricle. After injection, pups were placed on a 37 °C warming pad to recover from anesthesia and then were returned to the mother^92^.

### Statistical analysis

Statistical analysis was performed using GraphPad Prism version 8.0.0 (GraphPad Software, San Diego, California USA, www.graphpad.com), Matlab, and R software. Shapiro–Wilk test was used as a normality test, and the *F*-test was used to test for equal variances. Statistical comparison between two populations was performed using unpaired two-tailed Student’s *t*-test with Welch correction (when equal variance was not assumed) or Mann-Whitney *U* test non-parametric two-tailed test. For more than two normally distributed populations, the statistical comparison was performed with the one-way or two-way ANOVA with multiple comparisons or mixed-effects test with repeated measures. Statistical significance was considered at *p*-values < 0.05. Data are presented as mean ± SEM.

